# Proteogenomics analysis of human tissues using pangenomes

**DOI:** 10.1101/2024.05.24.595489

**Authors:** Dong Wang, Robbin Bouwmeester, Ping Zheng, Chengxin Dai, Aniel Sanchez, Kunxian Shu, Mingze Bai, Husen M. Umer, Yasset Perez-Riverol

## Abstract

The genomics landscape is evolving with the emergence of pangenomes, challenging the conventional single-reference genome model. The new human pangenome reference provides an extra dimension by incorporating variations observed in different human populations. However, the increasing use of pangenomes in human reference databases poses challenges for proteomics, which currently relies on UniProt canonical/isoform-based reference proteomics. Including more variant information in human proteomes, such as small and long open reading frames and pseudogenes, prompts the development of complex proteogenomics pipelines for analysis and validation. This study explores the advantages of pangenomes, particularly the human reference pangenome, on proteomics, and large-scale proteogenomics studies. We reanalyze two large human tissue datasets using the quantms workflow to identify novel peptides and variant proteins from the pangenome samples. Using three search engines SAGE, COMET, and MSGF+ followed by Percolator we analyzed 91,833,481 MS/MS spectra from more than 30 normal human tissues. We developed a robust deep-learning framework to validate the novel peptides based on DeepLC, MS2PIP and pyspectrumAI. The results yielded 170142 novel peptide spectrum matches, 4991 novel peptide sequences, and 3921 single amino acid variants, corresponding to 2367 genes across five population groups, demonstrating the effectiveness of our proteogenomics approach using the recent pangenome references.

## Introduction

The concept of pangenomes has recently transformed the traditional linear representation of a single reference genome ^1, 2, 3^. The Human Pangenome Reference Consortium has created a comprehensive and detailed human reference genome using a graph-based, telomere-to-telomere approach that aims to capture global genomic diversity ^3^. The human reference pangenomes are getting more attention with increasing support in ENSEMBL ^4^ and ENSEMBL Genomes ^5^. Human pangenomes and graph genomes represent a paradigm shift in genomics, moving away the field from singular reference genomes by accounting for the inherent genetic diversity of the human genome. Using long-read sequencing technologies, human reference pangenomes have improved the assembled contigs, especially in more difficult repeat elements. Pangenomes facilitate the comprehensive inclusion of genetic variants in reference assemblies which have an impact on transcriptomics and proteomics studies.

The advancements in the human reference pangenome provide an opportunity to improve the human reference proteome. The identification and quantification of peptides and proteins in mass spectrometry-based proteomics are usually performed with the UniProt reference proteomes that include canonical/isoforms protein sequences annotated by the UniProt Consortium ^6, 7^. Within UniProt, the concept of “canonical” is employed to denote the representative isoform of a protein, typically the one with the most complete and accurate sequence ^8^. The canonical isoform serves as a reference, aiding researchers in standardizing their analyses and interpretations of protein-related data within the context of the human proteome. However, the current canonical proteome database does not reflect the diverse genetic variations observed in the pangenome consortium. This has several implications, especially in identifying disease-associated protein variants.

To overcome this issue and identify proteins beyond the canonical proteome, researchers have incorporated genetic variants from public databases to identify variant peptides from MS data ^9, 10, 11, 12, 13^. While this approach has led to the identification of many variant peptides, its applicability is limited due to the large number of possible variants that lead to a substantial increase in the database size and thus the identification of many false-positive variant peptides. Also, studies aiming to identify disease-associated protein variants by incorporating variants observed only from the patients are hindered due to difficulty in distinguishing those that are due to the diversity in the human populations versus disease-specific peptide variants.

On the other hand, efforts to identify novel protein-coding regions beyond the canonical database have mainly focused on either direct or indirect in-silico translations of transcriptome and genome sequences to include intergenic regions, pseudo genes, long noncoding RNAs, etc. ^12, 14, 15^. This approach successfully identified thousands of novel peptides in regions not annotated in the canonical proteome. However, the database size issues and the lower resolution in the previous reference genome assemblies have impeded the identification of proteins, especially in contigs that were not well assembled due to DNA complexity and repeat elements in the previous human genome references. The new pangenome reference assembly which benefited from long-read technology provides a better chance to identify novel coding regions.

Proteogenomics workflows consist of three primary components: (i) a database of novel, non-canonical peptides, (ii) a search engine, analysis tool, or workflow facilitating the identification of the majority of novel evidence, and (iii) a validation framework designed to eliminate low-quality evidences (potentially false positives) from the novel discoveries ^13^. While traditional search engines such as MSFragger or MSGF+ are used to perform peptide identifications; multiple approaches are used to perform the validation and quality control of the novel peptide identifications ^13, 16, 17, 18^. Multiple tools have been developed to validate the variant peptides passing the FDR threshold, such as SAVControl ^19^, spectrumAI ^10^, and PepQuery ^20^. In addition, multiple authors have proposed the use of RT time prediction as a method for the validation of novel peptides ^11, 21^. However, the inflated FDR in proteogenomics studies remains a challenge while the development of novel algorithms and validation workflows remains an active topic within the field. The development of new and novel deep-learning tools that use the retention time and the spectra quality information in combination with existing tools could dramatically improve the detection of false positive non-canonical peptides in proteogenomics studies.

Here, we use protein annotations from the recent human pangenome project to reanalyze two large-scale human proteome datasets ^22, 23^. The *quantms* workflow ^24^ was used to perform parallel processing analysis of 2031 MS/MS files from more than 30 different tissues. quantms combines three different peptide identification tools SAGE ^25^, COMET ^26^, and MSGF+ ^27^ and uses the Percolator tool ^28^ to boost the number of peptide identifications, crucial in proteogenomics studies. Novel or non-canonical peptide identifications from pangenome proteins (called in this manuscript GCA peptides) were then validated using a robust deep-learning workflow. Our validation framework consists of two key components: a spectrum-based component and a retention time-based component. The spectrum-based component improves data quality by filtering out low-quality spectra based on the signal-to-noise ratio. It utilizes the MS2PIP algorithm ^29^ to compare predicted MS2 spectra with identified spectra, and the spectrumAI algorithm ^10^ to validate amino acid variants using b or y-ion evidence. We use predicted retention times from DeepLC ^30^ and SDRF annotations ^31^ to train a model with canonical peptides in the retention time-based component. This model is then used to predict retention times for novel peptides. In total, 4991 GCA peptides, and 3921 single amino acid variants were identified, corresponding to 2367 genes.

## Methods

### PXD010154: A complete map of healthy human tissues

The label-free dataset used in the manuscript is a comprehensive analysis of the proteome abundance of 29 healthy human tissues ^22^. In summary, the samples were collected from 13 male and 16 female healthy donors, and tryptic-peptide samples were analyzed in DDA mode generated using a Q-Exactive Plus mass spectrometer coupled online to a nanoflow LC system (NanoLC-Ultra). Peptides were fractionated via hydrophilic strong anion exchange (hSAX) offline chromatography, enabling deep tissue proteomic fractionation. The dataset was originally analyzed using ENSEMBL GRCh38 proteome using MaxQuant. We created a sample and data relationship format (SDRF) ^31^ file for the original dataset including the sample metadata, and data search settings including, for example, post-translational modifications, precursor and fragment tolerances (https://github.com/bigbio/pgt-pangenome/blob/main/PXD010154/PXD010154.sdrf.tsv).

### PXD016999: A Quantitative Proteome Map of the Human Body

Additionally, we used an isobaric (TMT) dataset, a quantitative map of the human body that includes data from different tissues ^23^. The study quantitatively profiled the proteome of 201 samples from 32 different tissue types of 14 healthy individuals. This was achieved using a tandem mass tag (TMT) 10plex/MS3 mass spectrometry strategy, which allows 10 isotopically labelled samples to be analyzed together. To improve proteome coverage, each sample was extensively fractionated. Tissue samples were randomized across TMT 10plex groups for cross-tissue comparison and to minimize technical variations between mass spectrometry runs. The SDRF of the given datasets was annotated and deposited in two different files depending on the instrument used: (https://github.com/bigbio/pgt-pangenome/blob/main/PXD016999/PXD016999-first-instrument.sdrf.tsv, https://github.com/bigbio/pgt-pangenome/blob/main/PXD016999/PXD016999-second-instrument.sdrf.tsv).

### Proteogenomics database construction

Protein-coding sequences from 97 samples were downloaded from the Pangenome project. The sequences were unified to obtain a single proteome across the samples (**Supplementary Tables - Table 1**). Additionally, reference human canonical proteins were obtained from ENSEMBL release 110. The resulting database contained 96,136 and 319,593 canonical and pangenome-derived proteins, respectively which translates to 782,775 and 892,614 unique tryptic peptides. A decoy database was generated using pypgatk ^12^ with the algorithm method decoy-pyrat ^32^.

### Large-scale identification workflow using quantms

We used the open-source workflow *quantms* ^24^ to perform the peptide and protein identification using the protein pangenome database. quantms is a nextflow workflow that allows for parallel/distributed proteomics data analysis using open-source tools, extensively benchmarked with the given datasets and UniProt canonical databases ^33, 34^. All the data was exported to the mzTab ^35^ file format and transformed into a quantms.io (https://github.com/bigbio/quantms.io) feature parquet file for fast and scalable data processing.

### Validation of novel peptides

Peptide spectrum identifications are first filtered by 1% global FDR using quantms workflow, in addition, all peptide matches need to be higher than 7 amino acids. Novel pangenome peptides (GCA peptides) are then aligned to common protein sequence databases (ENSEMBL GRCh38 - 122614 sequences, UniProt reviewed and unreviewed sequences with isoforms - 83975 sequences) to avoid reporting as novel peptides, miss-annotated matches.

To validate these novel peptides, we developed a workflow based on deep-learning tools DeepLC ^30^ and MS2PIP ^29^. In addition, we used spectrum-based properties such as signal-to-noise (SNR) for filtering low-quality spectra and the spectrumAI ^10^ algorithm (implemented in Python pyspectrumAI) which is specifically used to assess annotated ions flanking both sides of the substituted amino acid, to validate single amino acid variants (SAVs).

### Validation using MS2PIP

MS2PIP (MS/MS Peptide Ion Predictor) is a computational tool that predicts the fragment intensities of peptide tandem mass spectra (MS/MS). This means that the model estimates the intensity and the b and y ions for a given peptide sequence ^29, 36^. The use of MS2PIP allows for the calculation of the Pearson correlation between the predicted b and y ions and the observed spectrum identification. A higher correlation for b or y ions between the predicted and experimental spectrum indicates that the pipeline has made a better peptide identification.

Before applying MS2PIP we filter low-quality spectra using signal-to-noise ratio, removing MS2 spectra in the 5th and 95th percentile of the SNR distribution. Then, we computed the MS2PIP score as the combination of the b and y ions weighted Pearson correlations and the total number of ions predicted (Equation 1).

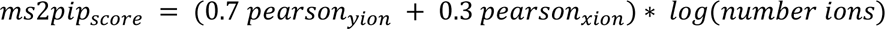

Similar to the signal-to-noise method for filtering, we leverage the distribution of all MS2PIP scores to determine the threshold at the 5th percentile, thereby eliminating peptide identifications with MS2PIP scores below 1.31 for dataset PXD010154 and 0.58 for PXD016999.

### Validation using spectrumAI

The spectrumAI algorithm was originally developed in R language by Zhu et al ^10^. spectrumAI is used to validate the single amino acid variant (SAAV) by trying to find the b or y-ions that provide evidence for the given variant. The algorithm first needs the non-canonical peptide to get aligned with the proteins to find the amino acid that changes. Both algorithms spectrumAI (pyspectrumAI) and the sequence alignment script were implemented in the package pypgatk (https://github.com/bigbio/py-pgatk) ^12^ enabling the user to detect the SAAV and validate it within the same Python package.

### Validation using DeepLC

DeepLC ^30^ is a deep-learning Python tool that enables the prediction of the retention time of modified peptides. DeepLC could be used with existing models from previous datasets or transfer learning using the same dataset’s experimental retention time. We adapted DeepLC version v2.2.27 (deeplc_novel.py in GitHub repository) to use the sample accession (source name) annotated in the SDRF and produce a model using the canonical peptides (GRCh38) and with the resulting model to predict the retention time of the GCA peptides. For each canonical peptide, the best posterior error probability (PEP) from all the peptide spectrum matches is selected. We calculate the difference between the predicted and experimental retention times for each novel peptide. We then calculate the relative error distribution and filter all the novel peptides in the 95th percentile. Additionally, the script visualizes error distributions and relationships between error percentiles and PEPs for analysis.

### Visualization and manual validation

Finally, all the resulting GCA novel peptides with their corresponding spectra are provided as USI (universal spectrum identifiers) ^37^ for manual validation (**Supplementary Tables - Table 2**). The PRIDE spectrum USI Viewer https://www.ebi.ac.uk/pride/archive/usi allows the user to visualize all the resulting pieces of evidence ^38^.

### Downstream and Network analysis

We developed a group of Jupyter notebooks and Python tools to perform the downstream analysis of the validated peptides, variants and genes (https://github.com/bigbio/pgt-pangenome). For the gene enrichment analysis, we developed a Jupyter notebook that uses the Gene Set Enrichment Analysis package GSEApy (https://github.com/zqfang/GSEApy) ^39^, and for different databases Reactome ^40^, GO Biological process, Molecular functions and Cellular components ^41^. The network analysis was performed using Cytoscape ^42^ and String database ^43^, and the selected genes were enriched using the complete human genome as the background. All peptides novel peptides were mapped to genome coordinates using the pepgenome tool (https://github.com/bigbio/pepgenome/) and the bed file was deposited in GitHub.

## Results

The pangenome project provides sequences of predicted proteins based on the assembly of each sample in the cohort of 97 individuals ^1, 2^. Although the combined number of predicted proteins is 3-fold higher compared to the number of proteins in the reference proteome (96,136 vs. 319,593), there is only a 12.3% increase in the number of tryptic peptides from the GCA samples indicating a large population diversity at the amino acid or peptide level (**Figure 1A**). Our goal here was to investigate the identification of these peptides at the proteome level. Due to the diversity of the samples and tissue types, this requires a wide search across multiple tissues and sample sources. To this end, we re-annotated the datasets PXD010154 and PXD016999 using SDRF ^31^. We analyzed the data using quantms ^24^ (**Figure 1B**) and three different search engines: COMET ^26^, MS-GF+ ^27^, and SAGE ^25^. The corresponding identified peptides were then boosted using the Percolator tool ^28^ and combined with ConsesusID ^44^. Finally, we applied 1% FDR at the peptide-spectrum match (PSM) level in both projects using the ProteomicsLFQ tool. The quantms workflow was run on the dataset PXD010154 (1367 MS files) and PXD016999 (664 MS files) at the EMBL-EBI compute cluster. Finally, a validation workflow for novel peptides was implemented to remove low-quality identifications (**Figure 1C**).

**Figure 1:**
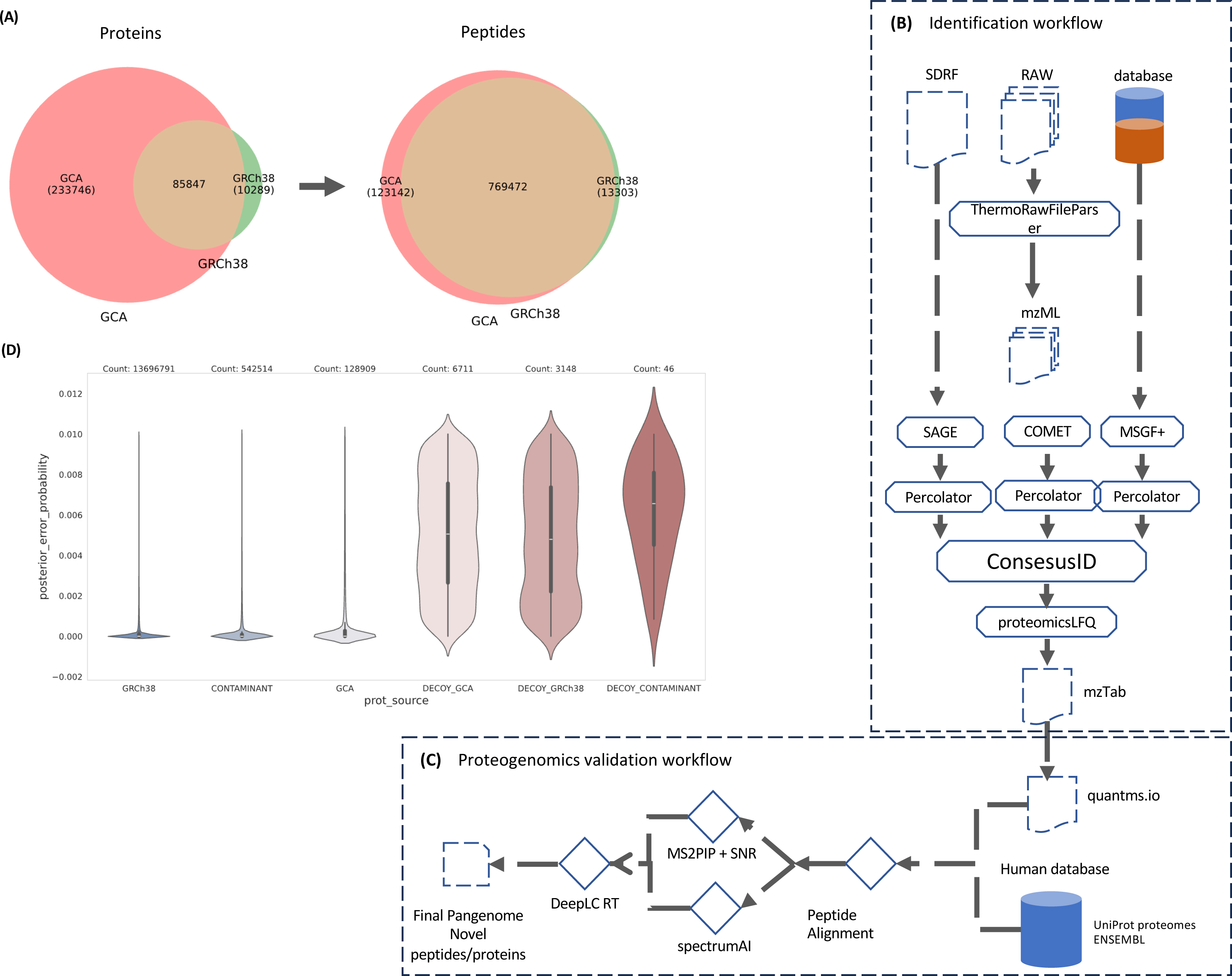
**(A)** Distribution of proteins and tryptic peptides for GCA pangenome and GRCh38 proteomes. **(B)** Identification workflow. Three different engines are used for peptide identification - SAGE, COMET and MSGF+; Percolator is then used to boost the number of identifications. The final output of the workflow is exported to mzTab format and quatms.io, a parquet file for data analysis. **(C)** Validation workflow including major steps, Peptide alignment of all peptides against major databases including UniProt Swiss-Prot and TrEMBL, and ENSEMBL GRCh38 protein database; the spectrumAI algorithm (implemented in the pyspectrumAI tool); the MS2PIP and Signal-to-Noise ratio step; and the final DeepLC (retention time prediction). **(D)** Distribution of posterior error probabilities for peptide spectrum matches (PSMs) from canonical and non-canonical pangenome peptides, contaminants, and decoys in dataset PXD010154.

We searched 2031 proteomic files (91,833,481 MS/MS spectra) from the PXD010154 (63,133,178 MS/MS) and PXD016999 (28,700,303) datasets covering more than 30 tissues. In dataset PXD010154, liver tissue was most represented with 144 spectra files followed by colon, lung, stomach, placenta and pituitary hypophysis each with 72 spectra files (**Supplementary Note 1 - Figure 1-2**). In the dataset PXD016999, the most represented tissue was the stomach with 263 MS files (**Supplementary Note 1 - Figure 3-4**). For data analysis on large PSM collections, we stored all PSMs in the fast and memory-efficient format quantms.io. quantms identified a total of 16,294,599 PSMs in dataset PXD010154, which corresponds to 508,840 peptide sequences; and 7,239,144 PSMs in PXD016999 corresponding to 285,330 peptide sequences. The distribution of PSMs by tissue for project PXD010154 is shown in **Supplementary Note 2 - Figure 5** whereas the tissue statistics for project PXD016999 are not shown since it is a TMT multiplex dataset thus each PSM represents multiple tissues through TMT channels. After applying a 1% FDR and aligning the PSMs with other proteome databases, including UniProt Swiss-Prot and TREMBL, we identified a total of 128,910 GCA PSMs in PXD010154 and 41,232 PSMs in PXD016999. The posterior error probability distribution of all PSMs is different for the three categories of target peptides: canonical, contaminants and non-canonical (**PXD010154 - Figure 1D, PXD016999 - Supplementary Note 3 - Figure 6**). While previous studies have applied class-specific FDR ^45, 46^, multiple studies have shown that additional validation is required ^10, 21^. In this study, instead of applying an extra FDR approach to filter for the novel peptides, we developed a workflow based on deep-learning tools; sample metadata, spectrum, and peptide properties to remove low-quality identifications (https://github.com/bigbio/quantms.io).

**Figure 2:**
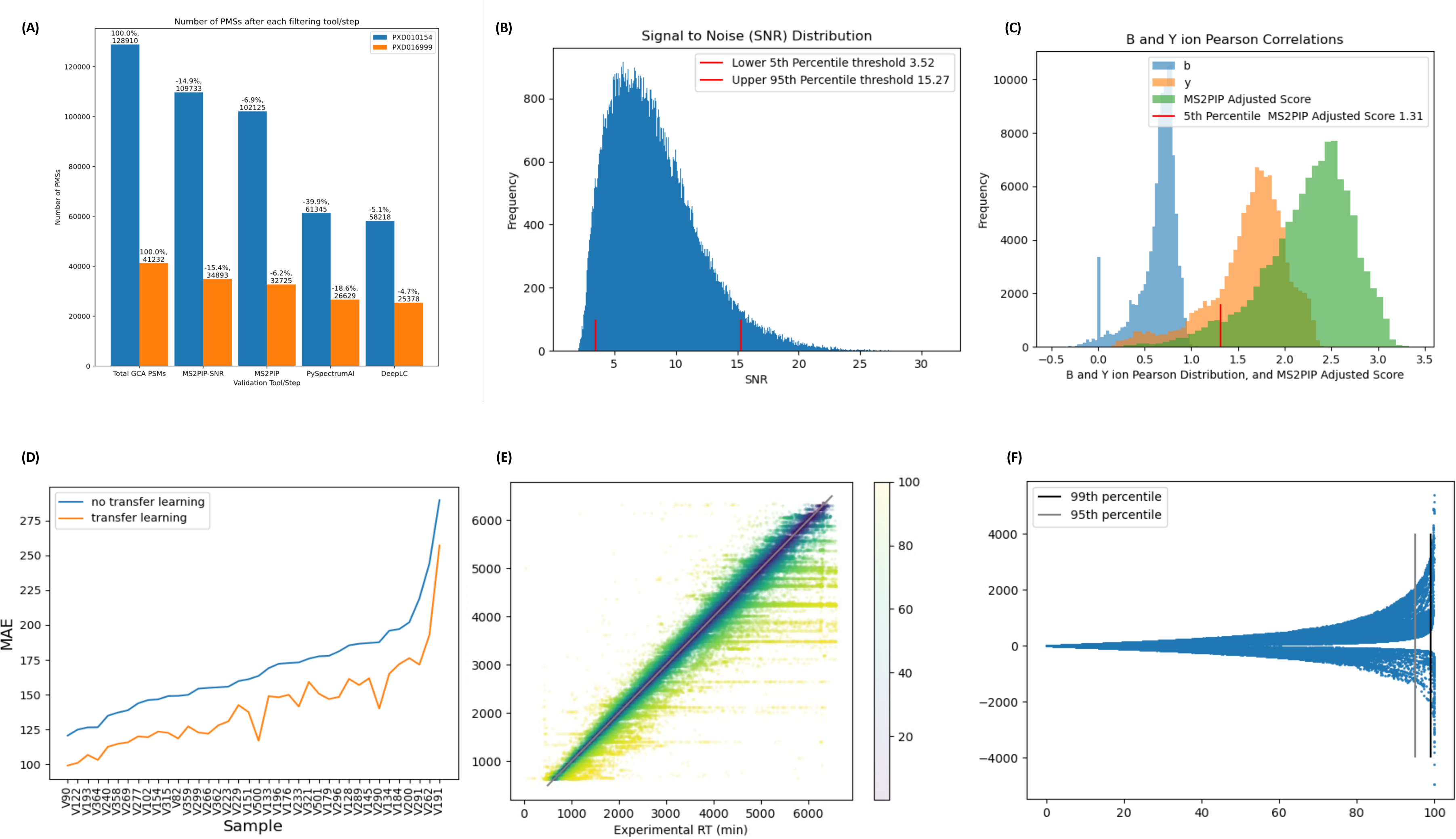
**(A)** Number and percentage of peptides filtered out **in** each step (MS2PIP + SNR; pyspectrumAI; DeepLC) of the validation pipeline. **(B)** Signal-to-noise ratio (SNR) distribution (SNR) for dataset PXD010154. In red the lower and upper threshold of the signal to noise that is used to remove low-quality peptide identifications. **(C)** MS2PIP score distribution for PXD010154; in blue and yellow the b and y-ion distributions; and in green the distribution for the adjusted score including the 5^th^ percentile (lower score) and threshold used to remove low-quality identifications. **(D)** Mean Absolute Error (MAE) for DeepLC transfer learning **versus** no transfer learning on PXD010154 **per** sample. **(E)** Correlation between experimental and predicted retention time for GCA peptides dataset PXD010154. **(F)** Relative Error distribution of DeepLC predictions for PXD010154 including the 99th and 95th percentile bars define peptides with the higher difference between predicted and experimental retention time.

**Figure 3:**
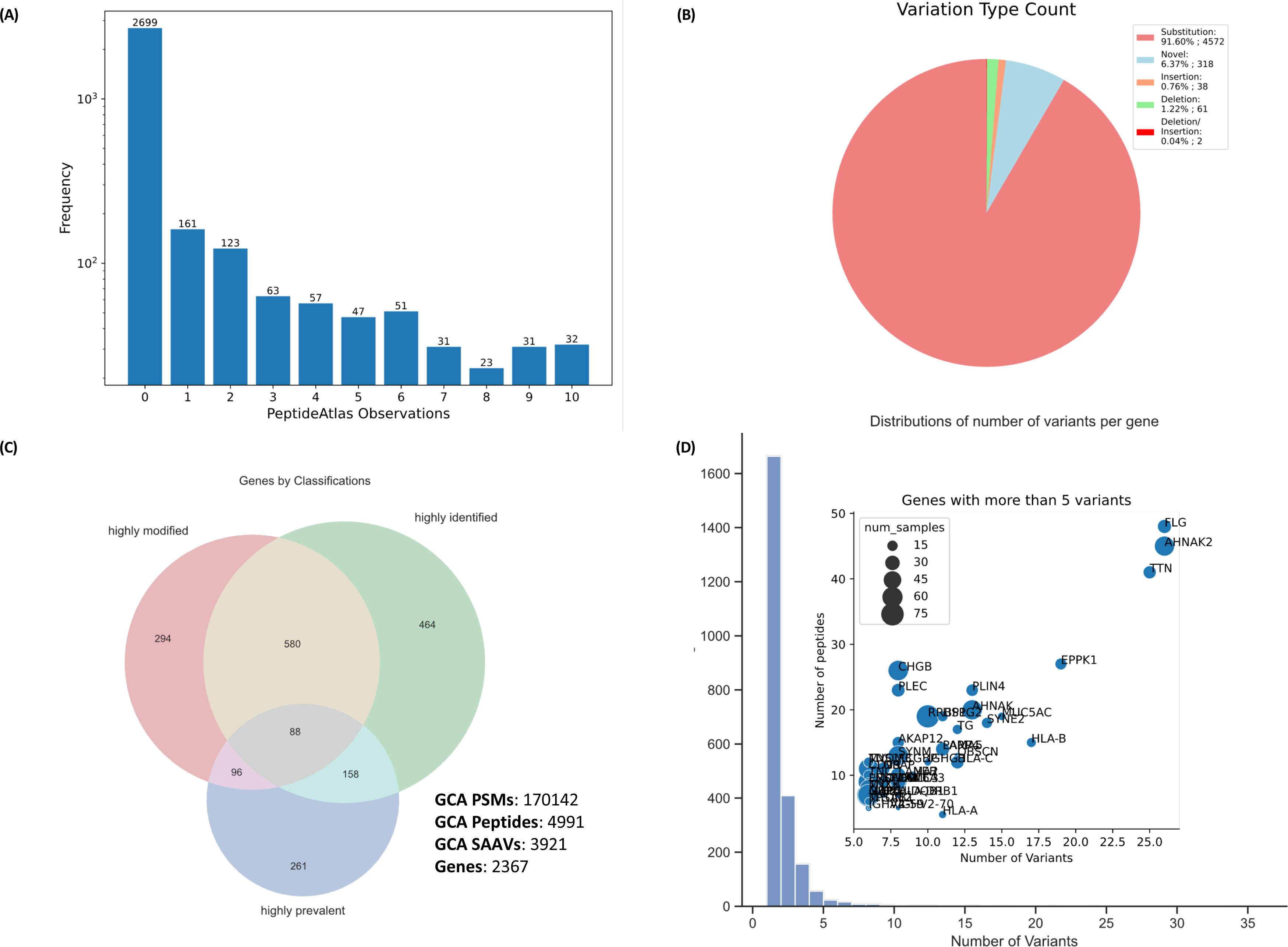
**(A)** Distribution of the number of observations for validated GCA peptides in the PeptideAtlas database. 2699 peptides have no previous observations in PeptideAtlas; 619 peptides have no more than 10 observations and 1673 have at least 11 observations. **(B)** Distribution of variant types for all final peptides. More than 91.6% of the GCA peptides result from single amino acid variants (SAAV), followed by novel peptides (peptides with more than one amino acid change from the canonical - 6.37%), and there were 38 insertions and 61 deletions and 2 that are both insertions and deletions. **(C)** Using a quantile analysis, we classify all genes into three categories: highly modified – number of variants; highly identified – number of peptides; highly presented – number of samples where the gene is observed. **(D)** Distributions of variants per gene and a scatter plot for each gene are plotted, with the number of variants, identified GCA peptides, and samples (size of the bubble) provided.

**Figure 4:**
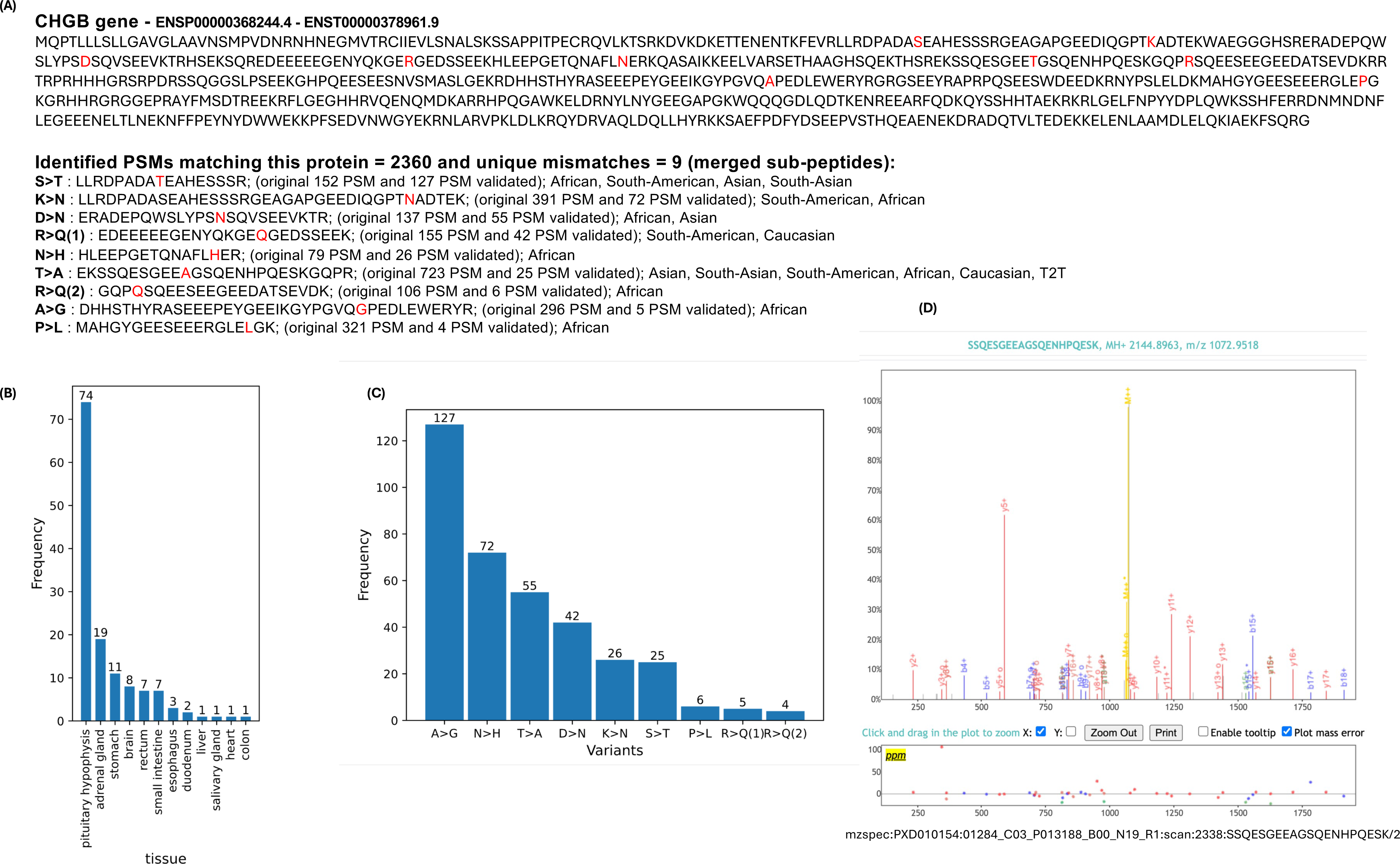
**(A)** Following the execution of the validation workflow for each variant site on the canonical protein CHGB (ENSP00000368244.4), including S>T, K>N, D>N, R>Q(1), N>H, T>A, R>Q(2), A>G and P>L, the average number of PSMs decreased from 262 to 40 per site. **(B)** Tissue distribution of PSMs at nine variant sites on the CHGB protein. The variants are mainly found in the pituitary hypophysis. **(C)** The number of PSMs at five variant sites on the CHGB protein, they are between 4 to 127 PMSs after the validation workflow. **(D)** Universal Spectrum Identifier (USI) for a validated PSM. Use PRIDE USI Viewer to view the visualization interface obtained by mzspec:PXD010154:01284_C03_P013188_B00_N19_R1:scan:2338:SSQESGEEAGSQENHP QESK/2

**Figure 5:**
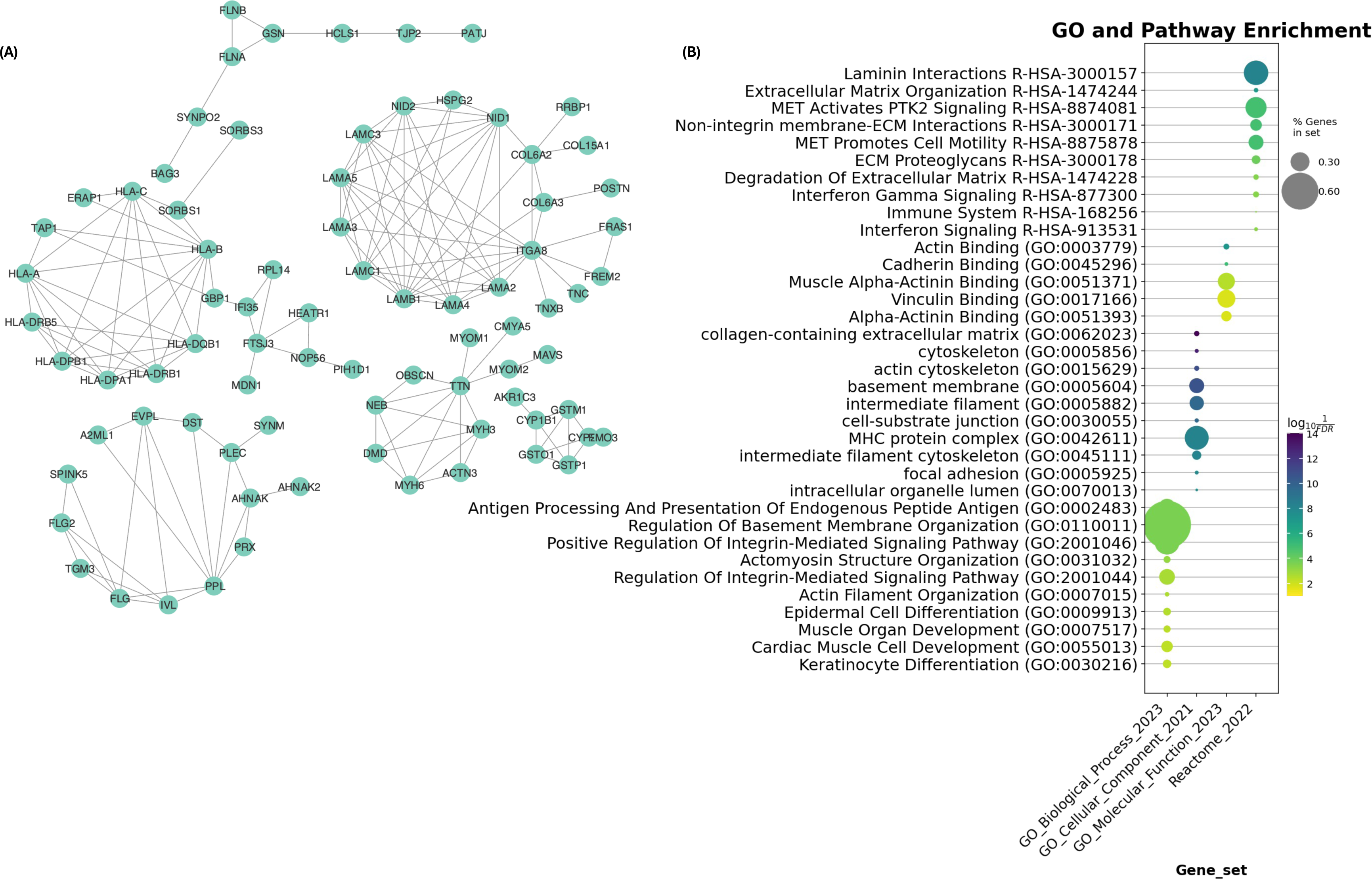
**(A)** The string network of most observed genes (higher number of peptides). **(B)** The functional enrichment of the network GO functional and biological processes and the Reactome pathways database, shows structural constituents of muscle, muscle structure development, muscle organ development, and muscle contraction have the most enriched functions, processes, and pathways.

**Figure 6:**
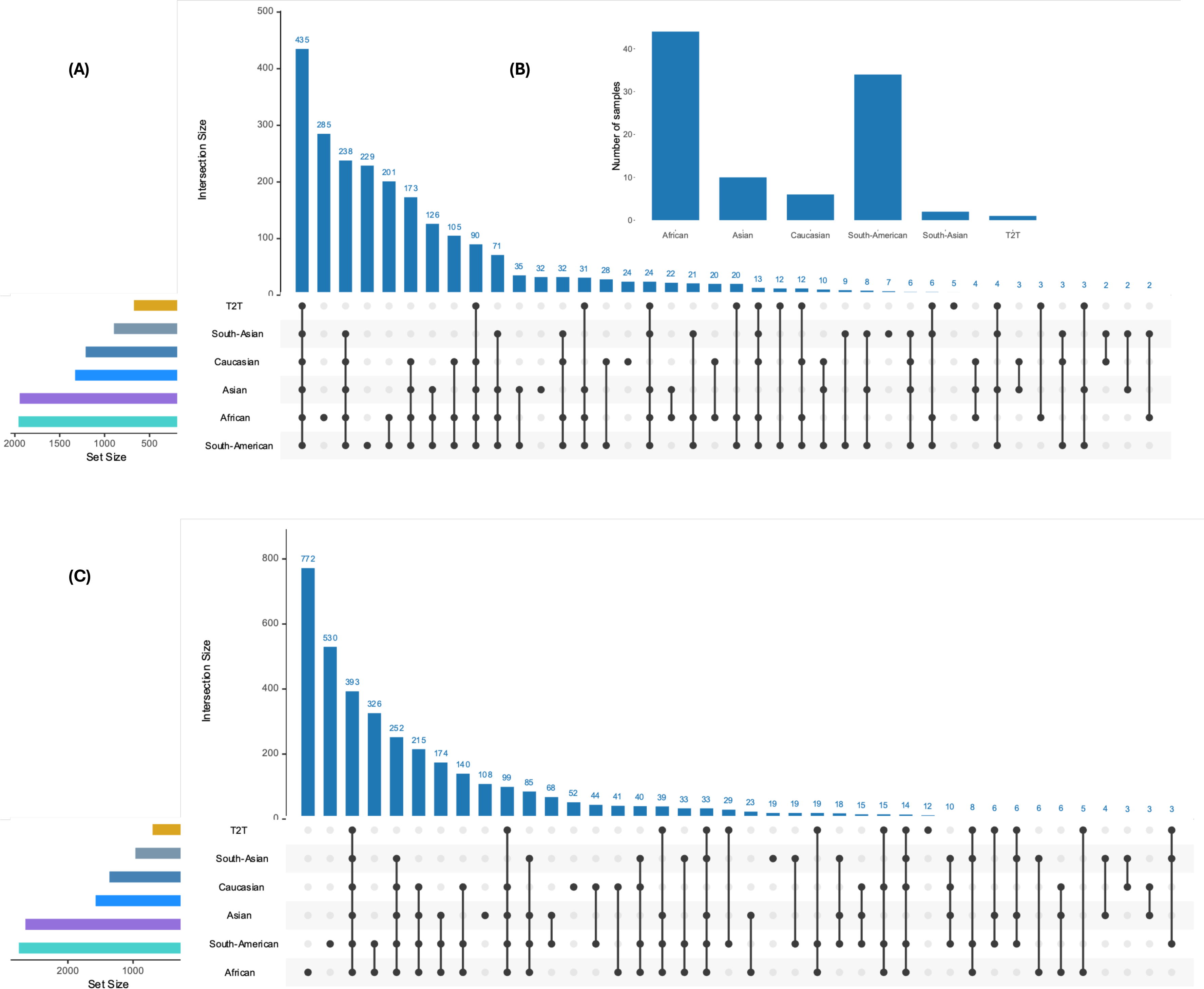
**(A)** Distribution of the number of genes identified with at least one variant and **(B)** variants by population group. **(C)** Number of samples in Pangenome by population group (African, Asian, Caucasian, South-American, South-Asian, T2T - Telomere-to-Telomere).

To this end, we applied four different algorithms consecutively (**Figure 2A**) to validate the peptide identifications and their corresponding spectra. First, we use a simple signal-to-noise ratio (SNR) algorithm to remove low-quality identifications supported by low-quality spectra. We use a distribution of the SNR scores and remove all the peptides in the lower 5th percentile or the 95th higher percentile (PXD010154 - **Figure 2B**, PXD016999 - **Supplementary Note 4 - Figure 7**). A total of 19177 PSMS (∼14.9%) in PXD010154 and 6339 PSMS (∼15.4%) in PXD016999 were removed by this simple signal-to-noise threshold eliminating low-quality spectra identifications (**Figure 2A**). Some removed peptide USIs are shown in **Supplementary Tables - Table 3.** Next, we use MS2PIP that predicts the spectrum for each peptide identification, followed by the calculation of Pearson correlation coefficients between the predicted *b* and *y* ions derived from both theoretical and experimental spectra. We applied the MS2PIP algorithm independent of the search engine to keep peptides with a higher number of *b* and *y* ions annotated (PXD010154 - **Figure 2C**, PXD016999 - **Supplementary Note 4 - Figure 8**). Different from previous approaches that use *b* and y ions Pearson correlations to filter peptides ^47^, we calculated a weighted score by combining the b and y ions scores and giving more weight to the y-ions. Using the weighted MS2PIP score a total of 6.9% and 6.2% of the GCA novel PSMs were removed (**Figure 2A**). Furthermore, in our spectrum-based validation framework, we used spectrumAI ^10^ to validate the corresponding variant amino acids. In this step ^10^we filtered out 39.9% and 18.6% peptide spectrum matches, for PXD010154 and PXD016999 respectively, with single amino acid variants (SAAV) that lacked *b* or *y* ion annotation validating the amino acid change (**Figure 2A**).

Finally, we used the DeepLC algorithm as an orthogonal validation step ^30^. Peptide retention time directly depends on the amino acid sequence and thus it can be used as an additional confirmation of identifications. Multiple studies have used retention time prediction to filter false-positive identifications for peptides with significant differences between the experimental and predicted retention time ^21, 47, 48^ since the predicted retention from the peptide sequence should be closer to the experimental retention; if the difference is high, the identified peptide is more likely to be a false positive. DeepLC could be used with existing models from previous datasets or by using the experimental retention from the same dataset via transfer learning. We benchmarked both approaches, using existing models (no transfer learning) and using the retention time reported for each canonical PSM on each sample (transfer learning) (**Figure 2D**). In our approach, we trained DeepLC based on the canonical peptide retention time (RT) and the sample metadata from SDRF and used the obtained model to predict the RT of GCA peptides and filter them. In each sample, transfer learning consistently resulted in a smaller mean absolute error (**Figure 2D**) and a higher Pearson correlation coefficient (**Supplementary Note 5 - Figure 9**). Our results show a higher correlation between experimental and theoretical retention time for GCA peptides (**Figure 2E**). We calculate the difference between the predicted and experimental retention times for each novel peptide. We then calculate the distribution of the relative error and filter all the novel peptides in the 95th percentile (**Figure 2F**).

Using DeepLC and retention time prediction, 5.1% and 4.7% of the GCA-matching PSMs were removed. A total of 83596 PSMs passed the four validation steps (**Figure 2A**) that corresponded to a total of 4991 peptide sequences, out of which 953 peptide sequences were identified in both projects. The 4991 GCA novel peptides were searched against the PeptideAtlas database ^49^ to investigate whether they have been identified in previous proteomics datasets since PeptideAtlas includes variant peptides in some of the projects that have utilized dbSNP ^50^ and other genomic variant sources (**Figure 3A**). Overall, 54.1% (2699) of the peptides have no previous observations in PeptideAtlas; 12.4% of the peptides (619) have fewer than 10 observations, whereas 33.5% (1673) have 11 or more observations ^50^. More than 91.6% of the GCA peptides result from single amino acid variants (SAAV), followed by novel peptides with 6.3% that have more than one amino acid change compared to the canonical protein. Additionally, there were 38 insertions and 61 deletions in addition to two peptides that were labelled as insertions and deletions depending on the matching protein source (**Figure 3B**). For more than 70% of these single amino acid variants, the counterpart canonical amino acid is identified at least once indicating large variation between samples.

### Sources of the GCA genes, and variants

Overall, 3921 SAAVs were identified in 2367 genes across all samples and tissues. The peptides were identified from proteomic datasets across various tissues (**Supplementary Tables – Table 4).** Ten tissues in PXD00154 were identified with more than 500 variant peptides (**Supplementary Note 6 - Figure 10)**; liver, lung, placenta, stomach, small intestine, and colon all identified at least 682 peptides, with their respective numbers being 1089, 936, 784, 767, 687, and 682. In comparison to other tissues, the pancreas had the fewest identified peptides, in total 189.

As expected, the majority of genes (94%) were associated with fewer than 4 variants whereas 138 genes had four or more variants (**Figure 3D**). Five genes had a very large number of AA variants: FLG, AHNAK2, TTN, EPPK1, HLA-B contained more than 20 variants each. Three of these genes (FLG, AHNAK2, and EPPK1) are highly expressed in skin tissues which may explain the high variation rates across populations (**Supplementary Note 7 – Figure 11**). Furthermore, HLA-B is known to encode many protein variants due to its role in the immune system. Interestingly, besides TTN, the number of GCA variant peptides identified per gene is not correlated with the protein length (**Supplementary Note 8 – Figure 12**). Noticeably, the variants in a single gene are identified in different samples and populations, for instance, gene CHGB contains 9 different single amino acid variants for the corresponding protein P05060 (**Figure 4A**). The variants are mainly found in the pituitary hypophysis in PXD010154 (**Figure 4B**) and the number of PSMs associated with each variant amino acid ranges between 4 to 127 PMSs after the validation workflow (**Figure 4C**). More importantly, after the validation workflow is run for each position, the number of PSMs drops on average from 262 PSMs to 40 PSMs (**Figure 4A**). Each validated PSM is supported by a universal spectrum identifier (USI) and can be visualized via PRIDE USI Viewer (**example Figure 4D**). We believe these large numbers of PSMs provide reliable evidence for the observed variations.

To summarize the genes with high variation rates, we conducted a quantile analysis to categorize genes based on the number of variants (highly modified), peptides supporting those variants (highly identified), and the number of GCA samples in which the genes are present (highly prevalent) (**Figure 3C**). Of 2367 genes, 294 are highly modified, 464 are highly identified, and 580 are highly prevalent in the pangenome samples. The combined total of these three groups is 911 genes. These genes are of significant interest, as their variants offer valuable insights beyond canonical genes typically used in proteomics studies. We further investigated the 294 highly modified genes that contained a large number of variants using the String network ^43^. Multiple highly connected networks are revealed across Laminin genes, including LAMC3, LAMA5, LAMA3, and LAMA; and HLA genes including HLA-A, HLA-B and HLA-C and HLA-DRB1; this aligns with expectations since HLA loci are known for their high polymorphism ^51, 52^. (**Figure 5A**) Interestingly, after performing enrichment analysis using the 294 genes; Twenty genes (TG, LAMA5, SYNE2, HLA-DRB1, AHNAK2, MKI67, FLG, HLA-C, PRUNE2, HLA-A, MUC16, HLA-B, HLA-DPB1, FRAS1, MUC12, OBSCN, TTN, FCGBP, NEB, PLIN4]) were previously reported by Vahdati and Wagner ^53^ as genes whose haplotype networks have significantly more squares in their networks than expected by chance alone (Table 1 – original manuscript ^53^). Additionally, Seventeen Genes in the network have been previously reported to be involved in the muscle cytoskeleton, and complex cytoskeletal networks that are critical for muscle contractile activity ^54^. The functional enrichment of these 258 genes, using GO ^41^ functional and biological processes; and the Reactome pathways database ^40^, shows Laminin interactions, regulation of basement membrane organization; and MHC protein complexes as the most enriched Reactome pathways; GO biological process; and GO cellular component (**Figure 5B**).

### Variation across populations

Only 1.66% (83) of the GCA peptides were identified in all pangenome samples and 17.99% (898) were sample-specific indicating a large sample specificity in protein variants. The remaining GCA peptides varied across the samples (**Supplementary Note 9 - Figure 13**). Overall, all samples had a large number of AA variants ranging between 545 to 703 supported by more than 1332 peptides per sample (**Supplementary Tables - Table 5**). Since the samples in the pangenome project are selected to represent global genetic diversity ^3^ we expect large variation in the identified variant peptides across the sample groups. To this end, we utilized the population information from the pangenomes project to group the samples according to their origin in addition to one sample group representing the telomere-to-telomere (T2T) genome reference.

Out of 2367 genes, 435 (17%) are found in all 6 major sample groups (South-Asian, Asian, Caucasian, African, South-American, and T2T samples). There were 285 and 229 genes identified specifically in African and South-American samples, respectively **(Figure 6A)**. We further explored the 285 variants that were observed only in the African population and identified by mass spectrometry using the genomAD database ^55^ (**Supplementary Note 10 - Figure 14**). Interestingly, more than 70% of the pangenome variants identified only in African samples and validated by mass spectrometry are also prevalent in other populations in genomAD, highlighting the capability of MS-based proteomics to detect widely frequent variants. For instance, we identified the variant 3-12817505-A-C (GRCh38) in gnomAD (https://gnomad.broadinstitute.org/variant/3-12817505-A-C?dataset=gnomad_r4), which corresponds to the SAAV variant p.His858Pro in the CAND2 gene and is common across all gnomAD populations. Additionally, our approach can identify variants with minimal prevalence in any gnomAD population, such as the variant 10-110881424-G-A (https://gnomad.broadinstitute.org/variant/10-110881424-G-A?dataset=gnomad_r4). These low-prevalence variants are particularly important because they are identified in the pangenome reference and validated by mass spectrometry in this study. This demonstrates the robustness of our proteomics analysis in uncovering both common and rare variants across diverse populations.

Overall, we identified the largest number of variant genes and peptides in the African and South-American population groups; almost four times higher than for other groups. However, this observation is unsurprising since these two populations had the highest representation in the pangenome project thus they had a larger contribution to our proteogenomics database (**Figure 6B**). Interestingly, at the variant peptide level, 393 variant peptides were identified in all populations **(Figure 6C)** whereas 1494 variant peptides (40%) were specific to one population only, suggesting that the identification and validation workflow can identify specific SAAVs in population studies, which has previously been challenging in proteomics ^22^.

## Discussion

The identification of a large number of single nucleotide variants (SNVs) highlights the importance of using pangenome references. Incorporating this diversity in proteomics studies allows for a more comprehensive understanding of protein variations across different populations. However, as multiple authors have previously highlighted ^13, 22, 45^, identifying protein SAAVs or novel coding sequences using proteomics is possible but demands specialized workflows and algorithms. We used three different search engines but also included Percolator to boost the number of identifications in proteogenomics studies. We performed an in-depth study using the SDRF ^31^ and quantms workflows ^24^ to analyze large-scale human tissue datasets with pangenome-based protein sequences. Instead of using a two-stage FDR approach, or class-FDR ^16, 56^; we combined multiple deep-learning tools and spectra properties to validate low-quality spectra. The signal-to-noise ratio (SNR) was used to remove peptides identified with low-quality spectra. Previous studies have shown that low-quality identifications have a higher signal-to-noise ratio, which makes them more likely to be false positives ^57, 58, 59^. In 2006, Nesvizhskii et al. ^59^ discussed the importance of using spectrum quality metrics such as signal-to-noise ratio (SNR) to improve peptide identification scores and detect false positives. While such metrics are not useful in proteomics studies based on canonical proteins and current high-resolution instruments, they are crucial in proteogenomics studies. This study found that SNR removes a large number (15%) of non-canonical low-quality identifications. Our SNR algorithm was essential in removing PSMs with both high and low signal-to-noise ratios as we observed that both extremes in the SNR distribution led to poor-quality identified spectra.

Additionally, MS2PIP algorithm was applied to ensure the identified peptides have a series of predicted b and y ions that are correlated with the experimental spectrum. We removed 6.8% of the identified PSMs using MS2PIP. These results are similar to a previous study by Levitsky et al. ^47^ that removed ∼5% of PSMs using a similar approach with the prediction tool PROSIT ^60^. Although SNR and M2PIP removed 21.8% of the low-quality PSMs, none of the algorithms validates the quality of a particular amino acid variant within a peptide identification. pyspectrumAI was the most stringent filtering step removing 34.8% of the peptides due to invalid b or y ion annotations at the AA variant sites. These results are similar to the findings reported in the original spectrumAI study ^10^ that showed b and y-ion annotations are essential to confirm the validity of variants in the specific positions. We applied a further step involving retention time and orthogonal spectrum validation to eliminate low-quality peptide-spectrum matches (PSMs) based on peptide amino acid conformation. Using the difference between accurately predicted peptide RT by DeepLC and observed RT as a quantitative metric, we managed to filter out 5% of possible false-positive identifications, which aligns with results reported in previous studies ^21, 48^. The main difference of earlier approaches is that we select the best canonical peptide identification for each sample or group of samples as indicated in the SDRF to train the model. This method helps create a model that improves the correlation between predicted and actual retention times for canonical and non-canonical peptides. Finally, as part of the validation framework developed in this study and following previous proteogenomics guidelines ^13^, we make all the USIs and spectra available for manual inspection in the PRIDE USI Viewer.

Our comprehensive analysis of two large-scale human tissue datasets identified a significant number of gene-coding variants across diverse samples, highlighting several key findings. We discovered 4,991 GCA peptides corresponding to 3921 single amino acid variants (SAAVs) in 2367 genes. The genes contained variants in multiple locations and samples, importantly all identifications were supported by strong evidence at the peptide level. The association of the highly modified, prevalent, and identified genes to known pathways and biological processes provide insights beyond the canonical genes typically used in proteomics, suggesting that incorporating these variant genes can enhance the accuracy and depth of proteomic studies ^61^. The functional enrichment analysis revealed significant networks and interactions within Laminin and HLA genes as well as MHC protein complexes, aligning with their known high polymorphism and roles in immune response and structural integrity. This finding, while expected, provides biological evidence of the accuracy of the developed identification and validation workflow in detecting single amino acid variants. Importantly, the variant peptides were identified across different tissues for which liver, lung, and placenta showed the highest counts, suggesting significant specific expression of the variant peptides. The variation observed across different population groups further supports the necessity of including diverse genetic backgrounds in pangenome references to capture the full spectrum of protein variants. For instance, only 1.66% of the GCA peptides were identified in all pangenome samples, while 17.99% were sample-specific, demonstrating a large sample specificity in protein variants. The African and South American populations had the largest representation in our proteogenomics database, and they had the largest number of population-specific variant peptides indicating the importance of conducting larger proteogenomics studies across larger populations to identify population-specific and common variants. Interestingly, 60% of the genes showed variation across all populations in our study emphasizing the wide-enriched coverage of variant amino acids detected using the pangenome. Our study demonstrates the significant potential of reusing public proteomics data combined with robust, large-scale proteogenomics workflows to identify and validate genome population variants by mass spectrometry.

## Code availability

The quantms workflow used to analyze the data is available at https://github.com/bigbio/quantms. The proteogenomics scripts and algorithms to generate the proteogenomics database, perform the peptide alignments and the pyspectrumAI are available in the pypgatk package https://github.com/bigbio/py-pgatk. Scripts and notebooks used for the downstream analysis are available at https://github.com/bigbio/pgt-pangenome.

## Data availability

The original RAW data for both datasets PXD010154 (https://www.ebi.ac.uk/pride/archive/projects/PXD010154), and PXD016999 (https://www.ebi.ac.uk/pride/archive/projects/PXD016999) can be found in PRIDE database. Results of the reanalysis using quantms and the given Pangenome database can be found at https://ftp.pride.ebi.ac.uk/pub/databases/pride/resources/proteomes/proteogenomics/tissues-pangenome/. Other complementary files and figures can be found at https://github.com/bigbio/pgt-pangenome.

## Contributions

Y. PR., H.M.U. conceived the project. Y. PR, H.M.U, D.W., R.B., C.D developed the workflow and analyzed the data. P.Z supported the team with the development of quantms.io for proteogenomics. A.S validated the spectra data and contributed to the development of the validation workflow. K. S., M. B. supported the students and contributed to the research discussions and funding. Y. PR., H.M.U and D.W mainly wrote the manuscript, although all authors contributed to the ideas and the content of it.

## Acknowledgements

Y.PR. acknowledge funding from Wellcome [grant number 223745/Z/21/Z], and EMBL core funding. D.W. and M.B. are supported by NSFC No.61501071.

## References

1. Gao Y, et al. A pangenome reference of 36 Chinese populations. Nature 619, 112–121 (2023).

2. Wang T, et al. The Human Pangenome Project: a global resource to map genomic diversity. Nature 604, 437–446 (2022).

3. Liao WW, et al. A draft human pangenome reference. Nature 617, 312–324 (2023).

4. Martin FJ, et al. Ensembl 2023. Nucleic Acids Res 51, D933–D941 (2023).

5. Yates AD, et al. Ensembl Genomes 2022: an expanding genome resource for non-vertebrates. Nucleic Acids Res 50, D996–D1003 (2022).

6. Bowler-Barnett EH, et al. UniProt and Mass Spectrometry-Based Proteomics-A 2-Way Working Relationship. Mol Cell Proteomics 22, 100591 (2023).

7. Aebersold R, Mann M. Mass-spectrometric exploration of proteome structure and function. Nature 537, 347–355 (2016).

8. UniProt C. UniProt: the Universal Protein Knowledgebase in 2023. Nucleic Acids Res 51, D523–D531 (2023).

9. Alfaro JA, Ignatchenko A, Ignatchenko V, Sinha A, Boutros PC, Kislinger T. Detecting protein variants by mass spectrometry: a comprehensive study in cancer cell-lines. Genome Med 9, 62 (2017).

10. Zhu Y, et al. Discovery of coding regions in the human genome by integrated proteogenomics analysis workflow. Nat Commun 9, 903 (2018).

11. Skiadopoulou D, Vasicek J, Kuznetsova K, Bouyssie D, Kall L, Vaudel M. Retention Time and Fragmentation Predictors Increase Confidence in Identification of Common Variant Peptides. J Proteome Res 22, 3190–3199 (2023).

12. Umer HM, et al. Generation of ENSEMBL-based proteogenomics databases boosts the identification of non-canonical peptides. Bioinformatics 38, 1470–1472 (2022).

13. Nesvizhskii AI. Proteogenomics: concepts, applications and computational strategies. Nat Methods 11, 1114–1125 (2014).

14. Zheng P, et al. Quantitative Proteomics Analysis Reveals Novel Insights into Mechanisms of Action of Long Noncoding RNA Hox Transcript Antisense Intergenic RNA (HOTAIR) in HeLa Cells. Mol Cell Proteomics 14, 1447–1463 (2015).

15. Olexiouk V, Crappe J, Verbruggen S, Verhegen K, Martens L, Menschaert G. sORFs.org: a repository of small ORFs identified by ribosome profiling. Nucleic Acids Res 44, D324–329 (2016).

16. Aggarwal S, Raj A, Kumar D, Dash D, Yadav AK. False discovery rate: the Achilles’ heel of proteogenomics. Brief Bioinform 23, (2022).

17. Tretter C, et al. Proteogenomic analysis reveals RNA as a source for tumor-agnostic neoantigen identification. Nat Commun 14, 4632 (2023).

18. Yao Z, et al. Proteogenomics of different urothelial bladder cancer stages reveals distinct molecular features for papillary cancer and carcinoma in situ. Nat Commun 14, 5670 (2023).

19. Yi X, Wang B, An Z, Gong F, Li J, Fu Y. Quality control of single amino acid variations detected by tandem mass spectrometry. J Proteomics 187, 144–151 (2018).

20. Wen B, Wang X, Zhang B. PepQuery enables fast, accurate, and convenient proteomic validation of novel genomic alterations. Genome Res 29, 485–493 (2019).

21. Wen B, Li K, Zhang Y, Zhang B. Cancer neoantigen prioritization through sensitive and reliable proteogenomics analysis. Nat Commun 11, 1759 (2020).

22. Wang D, et al. A deep proteome and transcriptome abundance atlas of 29 healthy human tissues. Mol Syst Biol 15, e8503 (2019).

23. Jiang L, et al. A Quantitative Proteome Map of the Human Body. Cell 183, 269–283 e219 (2020).

24. Dai C, et al. quantms: A cloud-based pipeline for proteomics reanalysis enables the quantification of 17521 proteins in 9,502 human samples. (2023).

25. Lazear MR. Sage: An Open-Source Tool for Fast Proteomics Searching and Quantification at Scale. J Proteome Res 22, 3652–3659 (2023).

26. Eng JK, Jahan TA, Hoopmann MR. Comet: an open-source MS/MS sequence database search tool. Proteomics 13, 22–24 (2013).

27. Kim S, Pevzner PA. MS-GF+ makes progress towards a universal database search tool for proteomics. Nat Commun 5, 5277 (2014).

28. The M, MacCoss MJ, Noble WS, Kall L. Fast and Accurate Protein False Discovery Rates on Large-Scale Proteomics Data Sets with Percolator 3.0. J Am Soc Mass Spectrom 27, 1719–1727 (2016).

29. Degroeve S, Martens L. MS2PIP: a tool for MS/MS peak intensity prediction. Bioinformatics 29, 3199–3203 (2013).

30. Bouwmeester R, Gabriels R, Hulstaert N, Martens L, Degroeve S. DeepLC can predict retention times for peptides that carry as-yet unseen modifications. Nat Methods 18, 1363–1369 (2021).

31. Dai C, et al. A proteomics sample metadata representation for multiomics integration and big data analysis. Nat Commun 12, 5854 (2021).

32. Wright JC, Choudhary JS. DecoyPyrat: Fast Non-redundant Hybrid Decoy Sequence Generation for Large Scale Proteomics. J Proteomics Bioinform 9, 176–180 (2016).

33. Bai M, Deng J, Dai C, Pfeuffer J, Sachsenberg T, Perez-Riverol Y. LFQ-Based Peptide and Protein Intensity Differential Expression Analysis. J Proteome Res 22, 2114–2123 (2023).

34. Wang H, et al. Tissue-based absolute quantification using large-scale TMT and LFQ experiments. Proteomics 23, e2300188 (2023).

35. Griss J, et al. The mzTab data exchange format: communicating mass-spectrometry-based proteomics and metabolomics experimental results to a wider audience. Mol Cell Proteomics 13, 2765–2775 (2014).

36. Declercq A, et al. Updated MS(2)PIP web server supports cutting-edge proteomics applications. Nucleic Acids Res 51, W338–W342 (2023).

37. Deutsch EW, et al. Universal Spectrum Identifier for mass spectra. Nat Methods 18, 768–770 (2021).

38. Perez-Riverol Y, et al. The PRIDE database resources in 2022: a hub for mass spectrometry-based proteomics evidences. Nucleic Acids Res 50, D543–D552 (2022).

39. Fang Z, Liu X, Peltz G. GSEApy: a comprehensive package for performing gene set enrichment analysis in Python. Bioinformatics 39, (2023).

40. Gillespie M, et al. The reactome pathway knowledgebase 2022. Nucleic Acids Res 50, D687–D692 (2022).

41. The Gene Ontology C. The Gene Ontology Resource: 20 years and still GOing strong. Nucleic Acids Res 47, D330-D338 (2019).

42. Shannon P, et al. Cytoscape: a software environment for integrated models of biomolecular interaction networks. Genome Res 13, 2498–2504 (2003).

43. Szklarczyk D, et al. The STRING database in 2023: protein-protein association networks and functional enrichment analyses for any sequenced genome of interest. Nucleic Acids Res 51, D638–D646 (2023).

44. Nahnsen S, Bertsch A, Rahnenfuhrer J, Nordheim A, Kohlbacher O. Probabilistic consensus scoring improves tandem mass spectrometry peptide identification. J Proteome Res 10, 3332–3343 (2011).

45. Karpova MA, et al. Exome-driven characterization of the cancer cell lines at the proteome level: the NCI-60 case study. J Proteome Res 13, 5551–5560 (2014).

46. Li J, et al. A bioinformatics workflow for variant peptide detection in shotgun proteomics. Mol Cell Proteomics 10, M110 006536 (2011).

47. Levitsky LI, et al. Massive Proteogenomic Reanalysis of Publicly Available Proteomic Datasets of Human Tissues in Search for Protein Recoding via Adenosine-to-Inosine RNA Editing. J Proteome Res 22, 1695–1711 (2023).

48. Li W, Wypych J, Duff RJ. Improved sequence variant analysis strategy by automated false positive removal. MAbs 9, 978–984 (2017).

49. Desiere F, et al. The PeptideAtlas project. Nucleic Acids Res 34, D655–658 (2006).

50. Sherry ST, et al. dbSNP: the NCBI database of genetic variation. Nucleic Acids Res 29, 308–311 (2001).

51. Barker DJ, et al. The IPD-IMGT/HLA Database. Nucleic Acids Res 51, D1053–D1060 (2023).

52. Palmer WH, Norman PJ. The impact of HLA polymorphism on herpesvirus infection and disease. Immunogenetics 75, 231–247 (2023).

53. A RV, Wagner A. Parallel or convergent evolution in human population genomic data revealed by genotype networks. BMC Evol Biol 16, 154 (2016).

54. Henderson CA, Gomez CG, Novak SM, Mi-Mi L, Gregorio CC. Overview of the Muscle Cytoskeleton. Compr Physiol 7, 891–944 (2017).

55. Chen S, et al. A genomic mutational constraint map using variation in 76,156 human genomes. Nature 625, 92–100 (2024).

56. Zhang K, et al. A note on the false discovery rate of novel peptides in proteogenomics. Bioinformatics 31, 3249–3253 (2015).

57. Xu H, Freitas MA. A dynamic noise level algorithm for spectral screening of peptide MS/MS spectra. BMC Bioinformatics 11, 436 (2010).

58. Castellana NE, et al. An automated proteogenomic method uses mass spectrometry to reveal novel genes in Zea mays. Mol Cell Proteomics 13, 157–167 (2014).

59. Nesvizhskii AI, et al. Dynamic spectrum quality assessment and iterative computational analysis of shotgun proteomic data: toward more efficient identification of post-translational modifications, sequence polymorphisms, and novel peptides. Mol Cell Proteomics 5, 652–670 (2006).

60. Gessulat S, et al. Prosit: proteome-wide prediction of peptide tandem mass spectra by deep learning. Nat Methods 16, 509–518 (2019).

61. Sinitcyn P, et al. Global detection of human variants and isoforms by deep proteome sequencing. Nat Biotechnol 41, 1776–1786 (2023).

