## Supplementary Notes for "Proteogenomics analysis of human tissues using pangenomes"

### **Supplementary Note 1**: Number of RAW files and MS/MS spectra by tissue for each dataset.

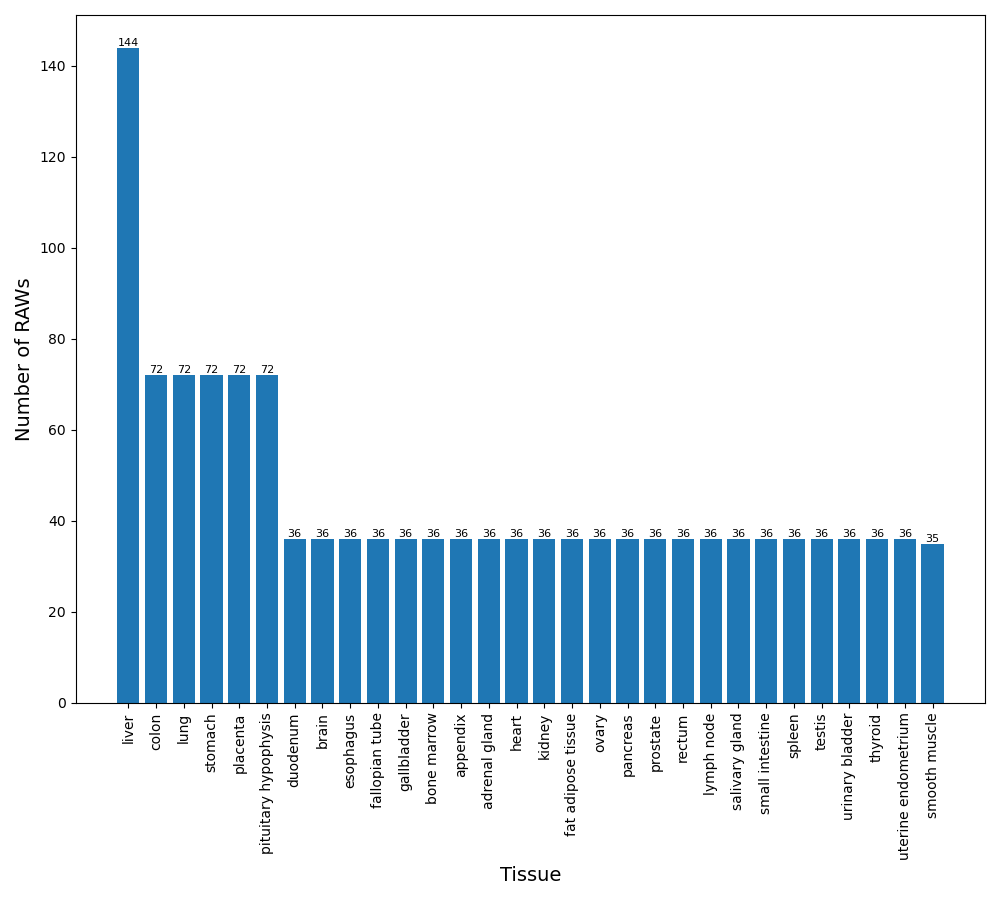

**Figure 1**: PXD010154 distribution of RAW files by tissue. The total number of RAW files is 1367, and the tissues are sorted by the number of files in the plot.

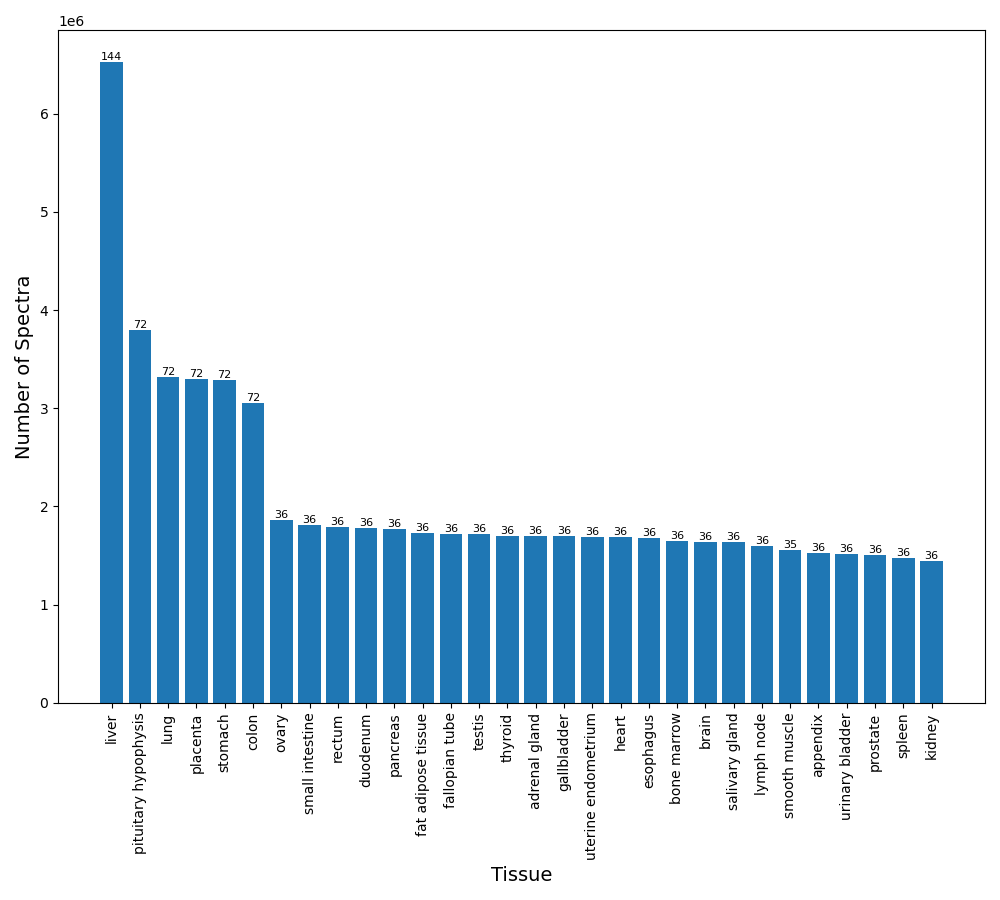

**Figure 2**: PXD010154 distribution of spectra (in millions) by tissue. The total number of MS/MS spectra analyzed is 63,133,178 and the number of MS/MS sorts the tissues in the plot.

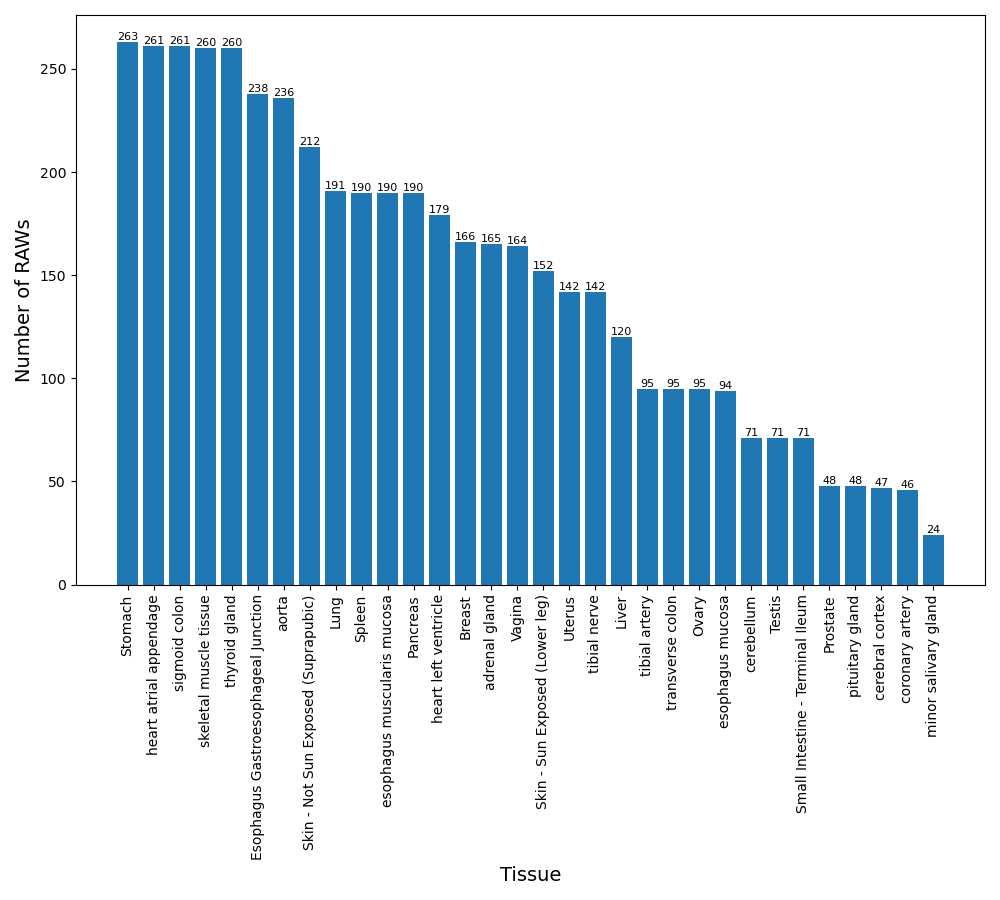

**Figure 3**: PXD016999 distribution of RAW files by tissue. The total number of RAW files is 664, and the tissues are sorted by the number of files in the plot.

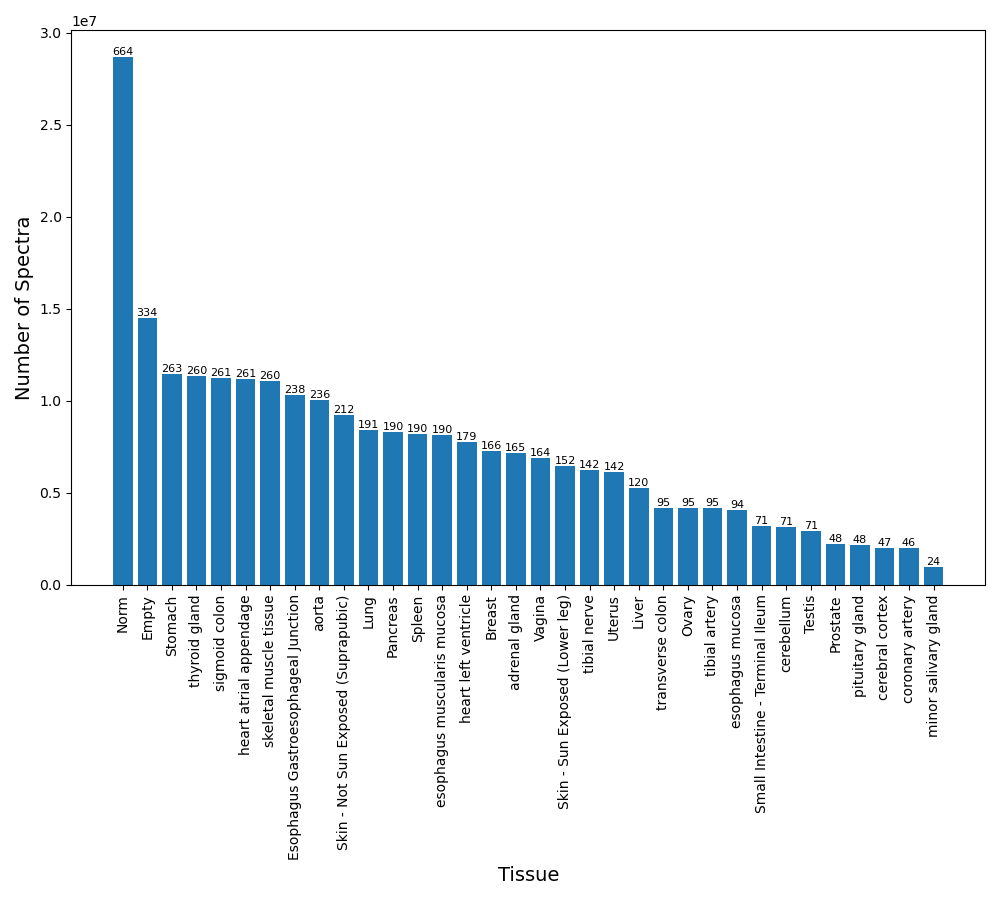

**Figure 4:** PXD016999 distribution of spectra (in millions) by tissue. The total number of MS/MS spectra analyzed is 28,700,303 and the number of MS/MS sorts the tissues in the plot. The Norm and Empty (tissues) are pooled channels used in the PXD016999 to increase the number of identifications.

### **Supplementary Note 2:** Distribution of PSMs by tissue for datasets PXD010154.

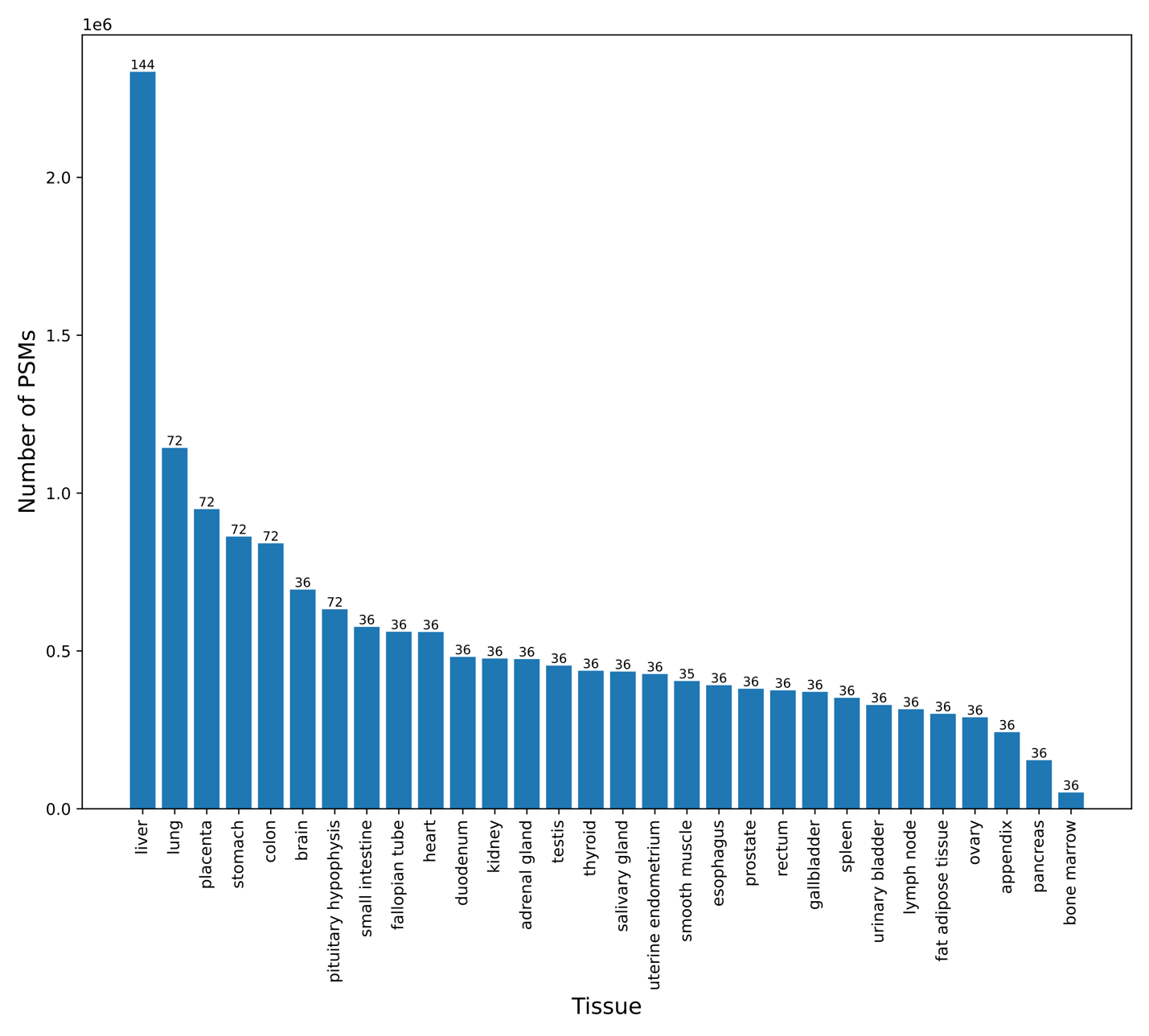

**Figure 5**: PXD010154 distribution of PSMs (in millions) by tissue. All tissues are sorted by the number of PSMs, which includes both canonical peptides (GRch38) and non-canonical peptides (GCA).

### **Supplementary Note 3:** PXD016999 distribution of PSMs and posterior error probabilities for the categories GCA, canonical, and contaminants.

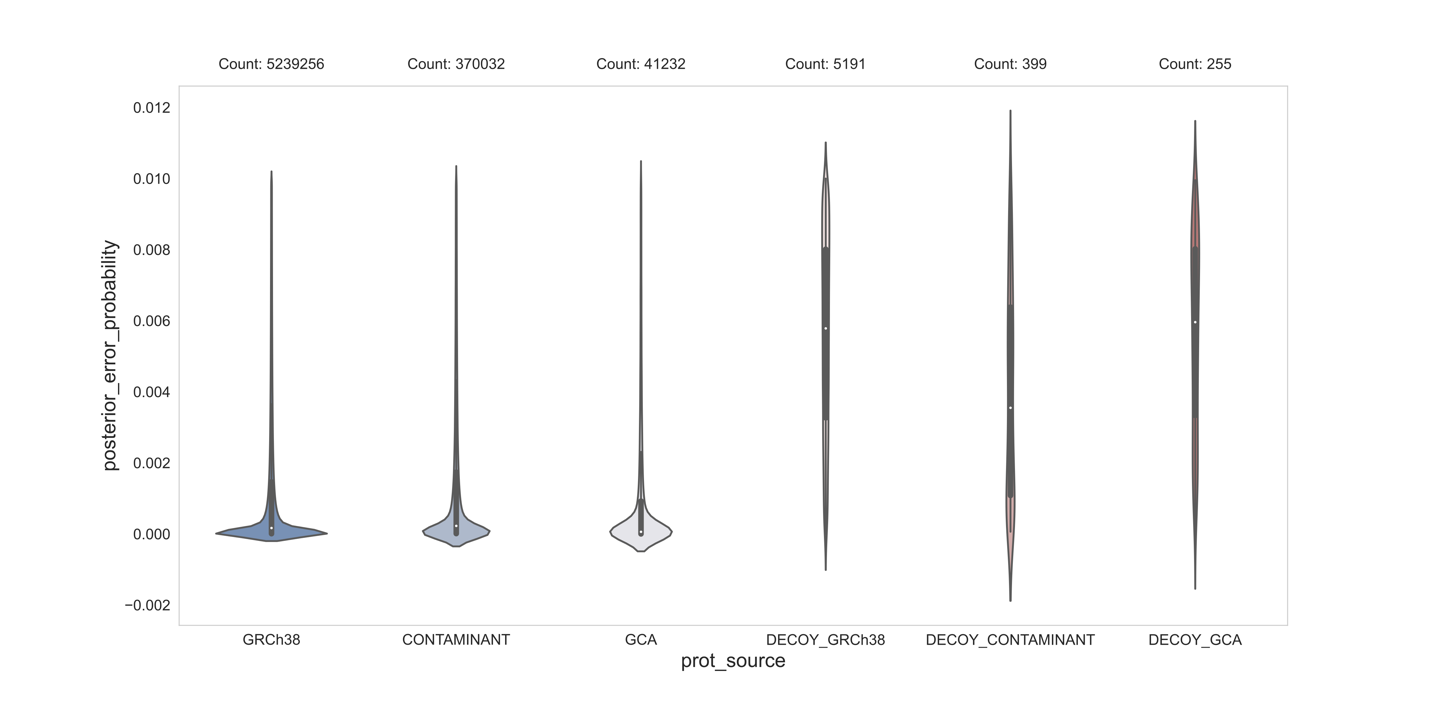

**Figure 6**: Violin plot with the distribution of the number of PSMs and posterior error probabilities for each category in dataset PXD016999, including canonical (GRCh38), contaminants, non-canonical (GCA), and their corresponding decoy counterparts.

### **Supplementary Note 4**: Spectra quality control results based on SNR and MS2PIP for PXD016999.

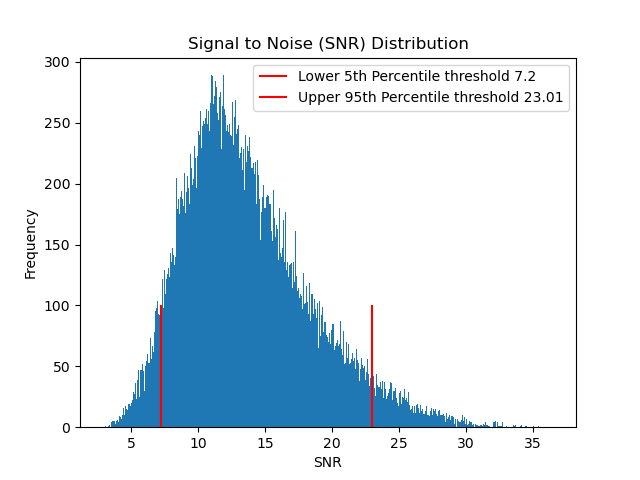

**Figure 7**: Signal-to-noise distribution for PXD016999 PSMs. The plot includes the line defining the 5th (7.2) and 95th (23.01) percentile threshold used to remove low-quality spectra.

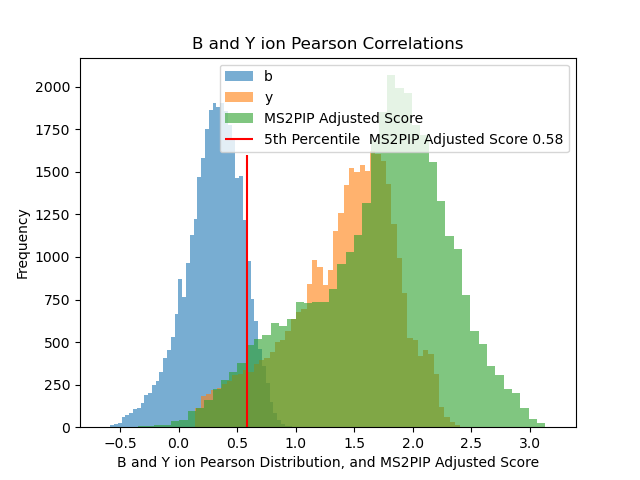

**Figure 8**: MS2PIP score distribution for PXD016999. The plot includes the threshold used to remove low-quality spectra (0.58).

### **Supplementary Note 5**: DeepLC results based on the training of canonical peptides and SDRF-proteomics metadata.

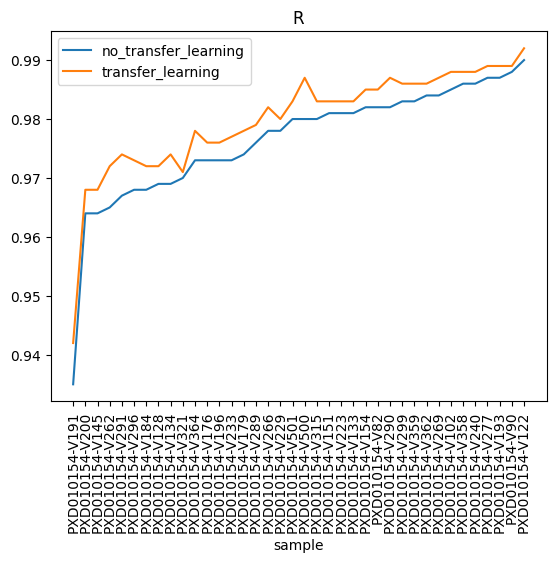

**Figure 9:** Pearson correlation between experimental and predicted retention time for every dataset sample PXD010154. The **orange** distribution is computed using transfer learning; where RT of the canonical peptides are used to train the model in DeepLC; and the **blue** distribution is the correlation between the experimental and predicted retention time using the default models in DeepLC.

### **Supplementary Note 6:** Number of identified GCA peptides per frequent tissue for PXD010154.

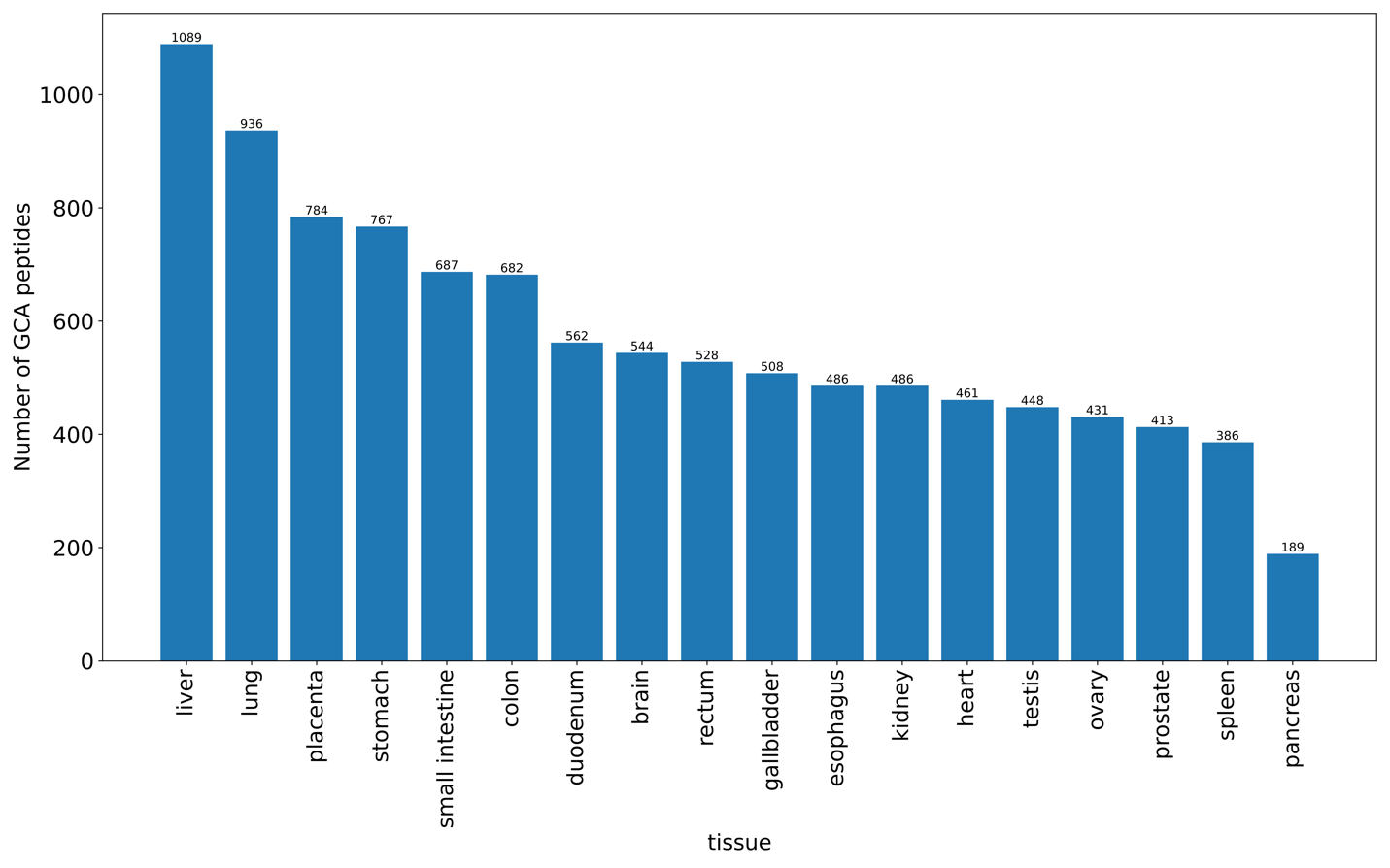

**Figure 10:** We selected tissues frequently studied in PXD010154 and counted the peptides identified in them. 10 of the 18 tissues identified at least 500 peptides, with liver, lung, placenta, stomach, small intestine and colon all identifying at least 682 peptides, and their numbers were 1089, 936, 784, 767, 687 and 682. Compared to other tissues, the pancreas had the fewest identified peptides, a total of 189.

**Supplementary Note 7:** Bulk tissue gene expression levels of FLG, EPPK1, and AHNAK2 based on GTEx.

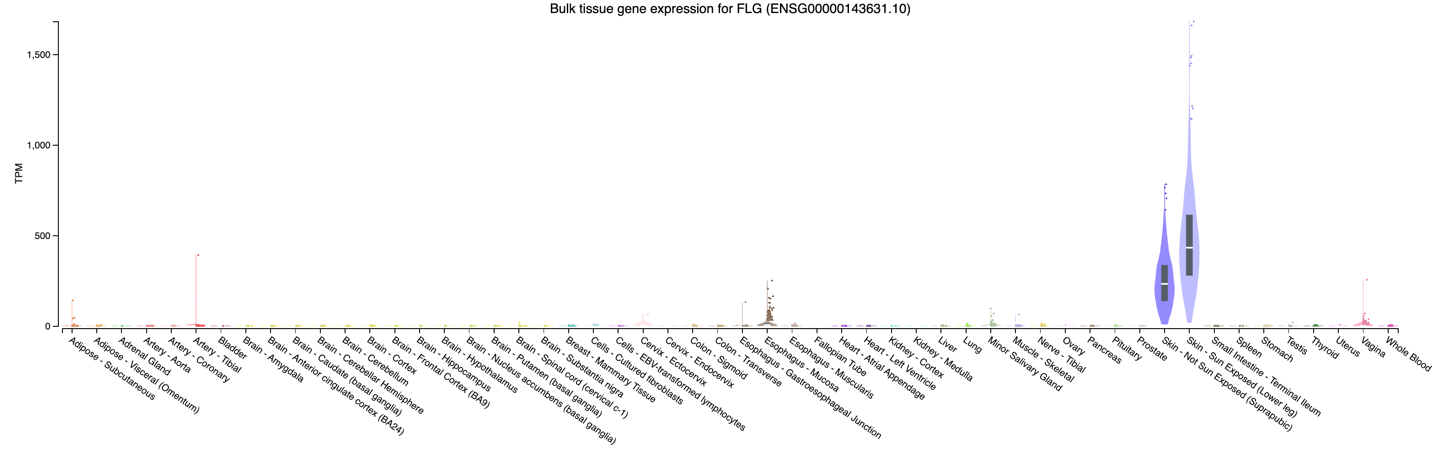

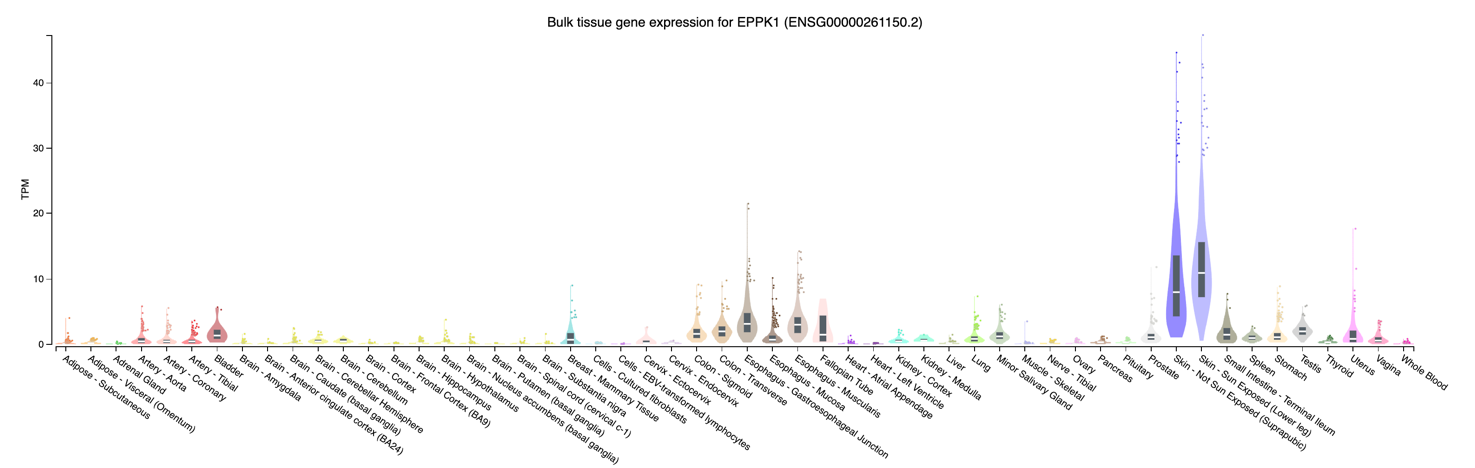

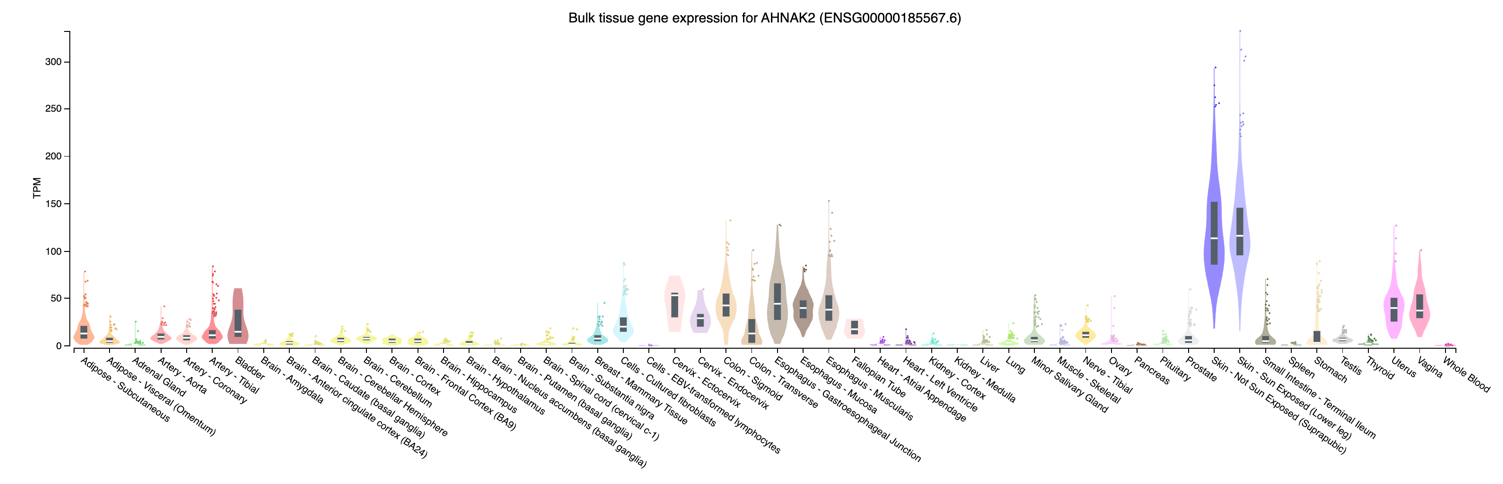

**Figure 11**: Bulk tissue gene expression levels of FLG, EPPK1, and AHNAK2 based on GTEx.

### **Supplementary Note 8**: Correlation between the number of GCA peptides identified per gene and Protein length.

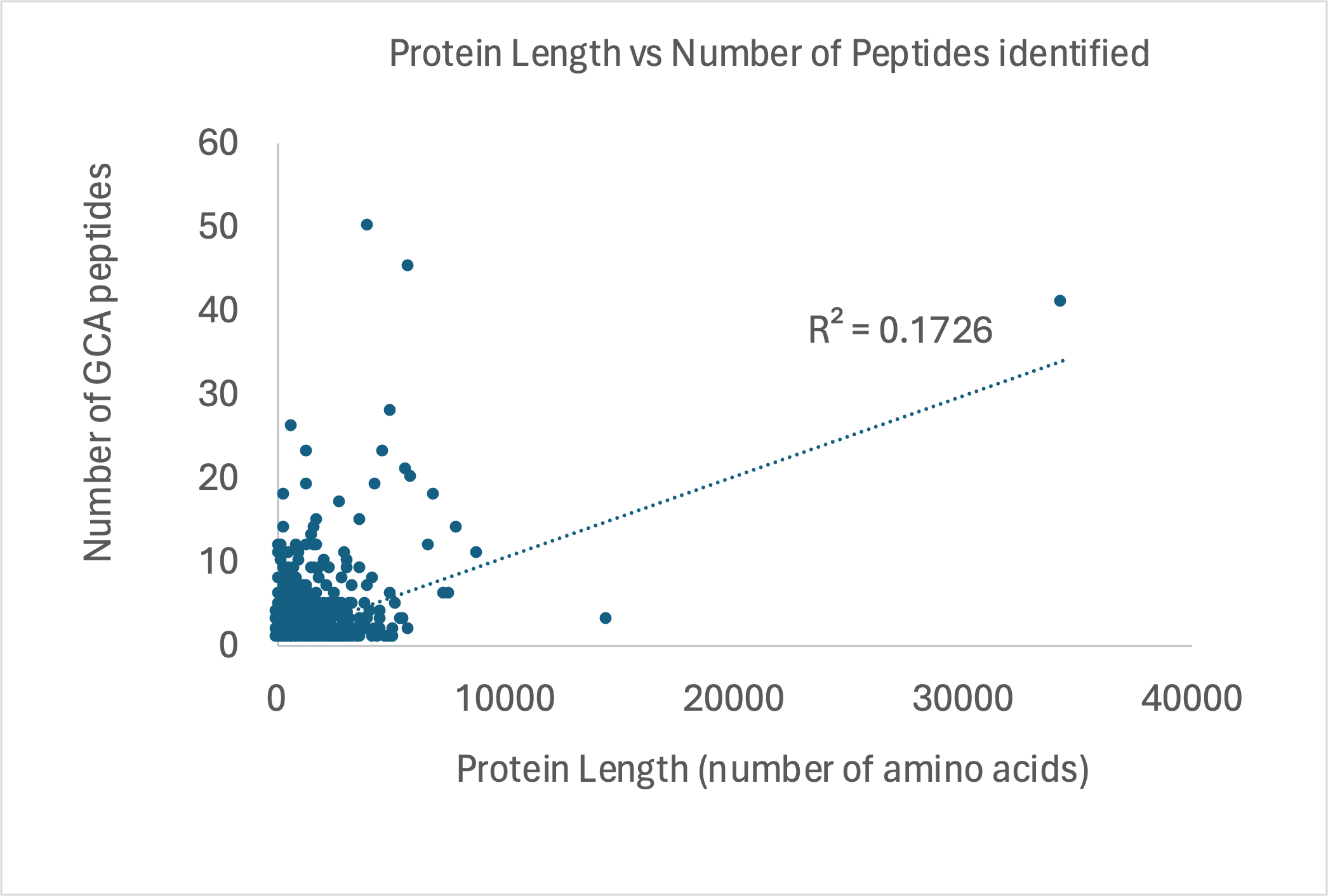

**Figure 12**: Correlation between the number of GCA peptides identified vs the length (number of amino acids) for the corresponding coding protein. We selected the protein canonical sequence to compute the protein length from the UniProt Swiss-Prot database. The Pearson correlation coefficient of 0.17 indicates that the number of novel GCA peptides identified is not correlated with the size of the proteins studied.

### **Supplementary Note 9:** Number of peptides identified in 97 GCA samples.

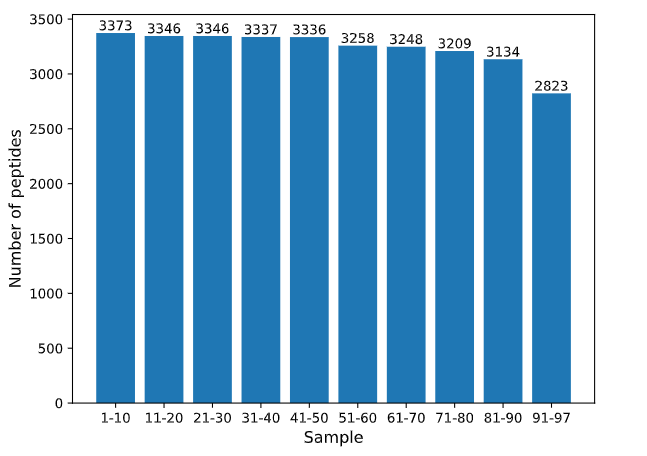

**Figure 13:** The number of peptides identified by the 97 GCA Samples from PXD010154 and PXD01699, divided into groups of 10.

### **Supplementary Note 10:** Variants distribution by population genomAD for those variants only identified in pangenome African samples.

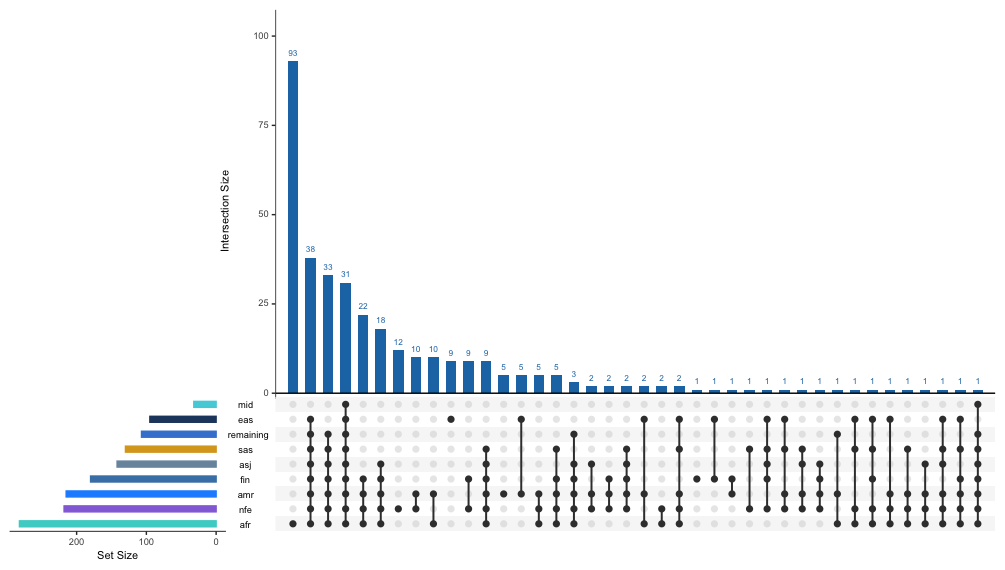

**Figure 14**: Distribution of variants shared across populations in genomAD. The variants under study are only the variants identified in the present proteogenomics study and only found in African samples in the pangenome. We filter the prevalence of a given variant in a population by using the RAW allele count in genomAD higher than 100. In the current plot, we used the code for each given population in genomAD: nfe - Non-Finnish European, amr - Latino/Admixed American, asj - Ashkenazi Jewish, afr - African/African American, mid - Middle Eastern, sas - South Asian, ami - Amish, eas - East Asian, remaining - Other or Remaining individuals (not specified), fin – Finnish.

### **Supplementary Note 11**: Population prevalence in genomAD for variant (SNV: 3-12817505 – A-C (GRCh38).

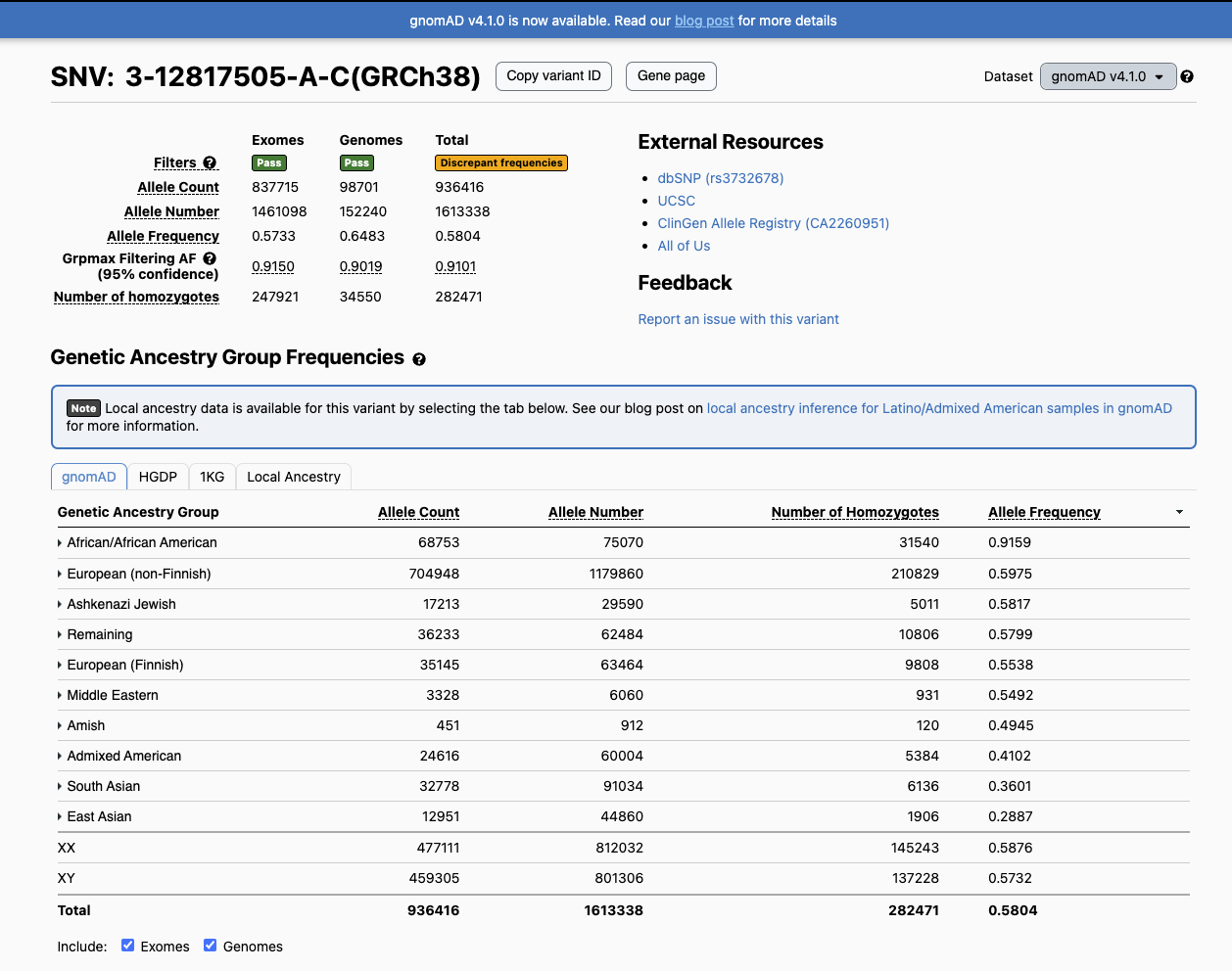

**Figure 15:** Prevalence of variant (SNV: 3-12817505 – A-C (GRCh38) - <https://gnomad.broadinstitute.org/variant/3-12817505-A-C?dataset=gnomad_r4>) in populations using the genomAD database. The variant corresponds to SAAV variant p.His858Pro in gene CAND2 (<https://gnomad.broadinstitute.org/region/3-12817452-12817512?dataset=gnomad_r4>)
